# Transcriptomic profiles of single-copy marker genes enable predicting bacterial growth states in microbial communities

**DOI:** 10.1101/2025.08.26.672432

**Authors:** M. L. Staeubli, H. J. Ruscheweyh, G. Greter, M. Dmitrijeva, S. Miravet-Verde, M. Arnoldini, W. D. Hardt, E. Slack, A. Sintsova, S. Sunagawa

**Affiliations:** Institute of Microbiology, Department of Biology, ETH Zurich, 8093 Zurich, Switzerland and Swiss Institute of Bioinformatics; Institute for Food Sciences, Health and Nutrition, Department of Health Sciences and Technology, ETH Zurich, 8093 Zurich, Switzerland; Institute of Microbiology, Department of Biology, ETH Zurich, 8093 Zurich, Switzerland; Sir William Dunn School of Pathology, University of Oxford, United Kingdom; Basel Research Center for Child Health, Switzerland

## Abstract

Studying microbial community dynamics is fundamental to better understand ecosystem stability, resilience and environmental change. Community composition changes with the growth of individual members, yet current methods to estimate microbial growth in communities face substantial limitations. For example, genome sequence-based estimates of maximum growth rates may not reflect growth patterns in the natural environment well, and metagenomic *in situ* growth prediction requires the availability of reference genomes and shows limited accuracy for slow-growing bacteria. Gene expression data provide an information-rich readout of community activity that could reflect growth, however, cross-species comparisons in community settings remain challenging. An approach using expression signatures of universal, single-copy marker genes provides independence from reference genomes and may thereby enable comparability across species. Here, we present a transcriptomic, marker gene-based growth classifier that predicts the growth states of bacterial strains from different phyla cultivated in diverse conditions. We demonstrate its application *in vivo* in gnotobiotic mice carrying the same bacterial strains, and in a more complex synthetic community, where predicted growth states align with reported growth inhibition induced by systemic inflammatory response. This approach offers a new method for predicting bacterial growth states across species, with potential for broad application in the study of microbial growth dynamics at the whole community level.

## Introduction

In natural environments, diverse microbes form complex communities that carry out important functions affecting host and ecosystem health. Depending on environmental conditions, microbial community composition changes as a function of growth, death and migration. Studying these dynamics is central to better understanding ecosystem stability and resilience upon environmental perturbations^1–3^. For example, microbial community dynamics can alter nutrient cycling and carbon storage in soils^4^, impact primary productivity and biogeochemical cycles in the ocean^5–7^ and modulate disease susceptibility of plants and animals^8,9^.

Despite its importance, measuring, modeling and understanding microbial community dynamics remains a major challenge^10–13^ with different approaches showing specific limitations. While metagenomic sequencing has facilitated assessing community composition at large scale^14–16^, deciphering growth patterns of individual community members has remained elusive. RNA:DNA ratios of the 16S ribosomal RNA gene have been used to distinguish activity and dormancy^17^, however these ratios may represent activity rather than growth, depend on gene copy number and rely on arbitrary thresholds, limiting their generality across taxa^18^. Similarly, genome-derived metrics, such as codon usage biases^19–22^ or rRNA operon copy numbers^23^ have been used to predict maximum growth rates, but these are rarely reached in natural environments^24^ and thus provide limited insight on growth *in situ*.

Aimed at predicting bacterial growth rates *in situ*, recent methods use whole-genome shotgun sequencing of microbial community DNA (metagenomes) and align the resulting reads to reference genomes^25–29^. These methods assume circular genomes, bi-directional DNA replication from one origin, and a direct relationship between the time needed for replication and cell division. They leverage the decreasing sequencing coverage from the origin of replication (*ori*) to the terminus to infer Peak-to-Trough Ratio (PTR) estimates as a proxy for *in situ* growth rates. However, despite the conceptual simplicity, these methods have also shown limited applicability, accuracy and interpretability. First, the varying quality and availability of reference genomes may bias PTR estimates or altogether prevent generating estimates for certain taxa, especially in currently underexplored environments^30^. Secondly, there are exceptions to the assumptions (e.g., linear chromosomes, multiple replication origins or chromosomes, asymmetric division), that may lead to different relationships between genome sequencing coverage and growth rates^31–34^. Lastly, log_2_(PTR) estimates for marine metagenomes suggest low accuracy for slow-growing bacteria^35^, which may represent the majority of community members, particularly in nutrient-scarce environments. Altogether, these limitations prevent obtaining reliable *in situ* estimates to study microbial growth dynamics at the whole community level.

Alternatively, transcriptomic data may be leveraged to study gene expression signatures informative of microbial activity or growth. Global transcriptome shifts across growth phases *in vitro* have revealed direct relationships between the requirements for ribosomes and growth^36^. However, finding and comparing growth-associated transcriptomic signatures across species and in community settings remains challenging^37^. Thus far, transcriptomic signatures have not been investigated with the objective to develop a universally applicable method for growth prediction. While one recent study^38^ predicted growth rates based on gene expression in *E. coli*, the developed model depends on the availability of complete reference genomes and thus would not be applicable to other species. Testing the transferability of transcriptomic-based growth prediction across diverse bacterial species would require the use of a shared set of genes, ideally present in one copy per genome. Such universal single-copy marker genes (MGs) have been used for prokaryotic species delineation^39^, phylogenetic tree reconstruction^40^, and taxonomic profiling of microbial communities^30,41^, while their suitability for predicting microbial growth has not been assessed so far. These MGs show promise for transcriptomic growth predictions because they contain many ribosomal protein genes and other genes involved in translation^42^.

In this work, we generated paired genomes and transcriptomes of three bacterial strains from different phyla that were grown under diverse conditions. We analyzed global gene expression changes across growth phases and evaluated the use of transcriptomic MG profiles for growth prediction. To this end, we developed a binary classifier predicting bacteria as growing or non-growing according to the similarity of transcriptomic MG profiles to exponential or stationary phase, respectively. To evaluate the classifier in a community setting *in vivo*, we applied it to cecum content samples of gnotobiotic mice containing the same three strains. To evaluate its applicability beyond these strains, we classified the growth states of strains from three other species previously suggested to experience growth inhibition through lipopolysaccharide-induced systemic inflammatory response^43^. The proposed method complements PTR-based approaches by i) solely depending on marker genes rather than complete reference genomes, ii) increasing interpretability through providing confidence score-based growth classifications and iii) demonstrating *in vivo* applicability including for slow-growing bacteria.

## Results

### Growth phase-dependent transcriptomic signatures in bacterial strains from different phyla

To enable the study of transcriptomic signatures of bacterial growth, we cultivated three strains from different phyla (*Escherichia coli*: Pseudomonadota, *Bacteroides thetaiotaomicron*: Bacteroidota and *Agathobacter rectalis*: Bacillota) with varying carbon sources, pH and temperature (Fig 1a; Supplementary Table 1). We collected samples at multiple time points across growth phases and estimated growth rates from changes in optical density over time (ΔOD). Exponential phase (EX) samples were collected during log-scaled linear increase in OD and stationary phase (ST) samples were collected at stable OD with net zero growth (ΔOD ≈ 0) after carrying capacity was reached. In *E. coli*, up to 3 consecutive transition phase (TR) samples were collected in between EX and ST phase from a slower increase in OD, temporary first plateau or slow decrease. Depending on the strain and culture conditions, the exponential growth rates of *E. coli* (0.10 to 0.94 h^-1^) and *B. theta* (0.12 to 0.76 h^-1^) were similarly fast, while *A. rectalis* was growing at considerably slower rates (0.02 to 0.23 h^-1^) (Fig. 1b). Co-extracting and shotgun sequencing of DNA and RNA from these samples resulted in 257 paired genomes and transcriptomes, allowing us to analyze genomic and transcriptomic signatures of bacterial growth.

**Figure 1.**
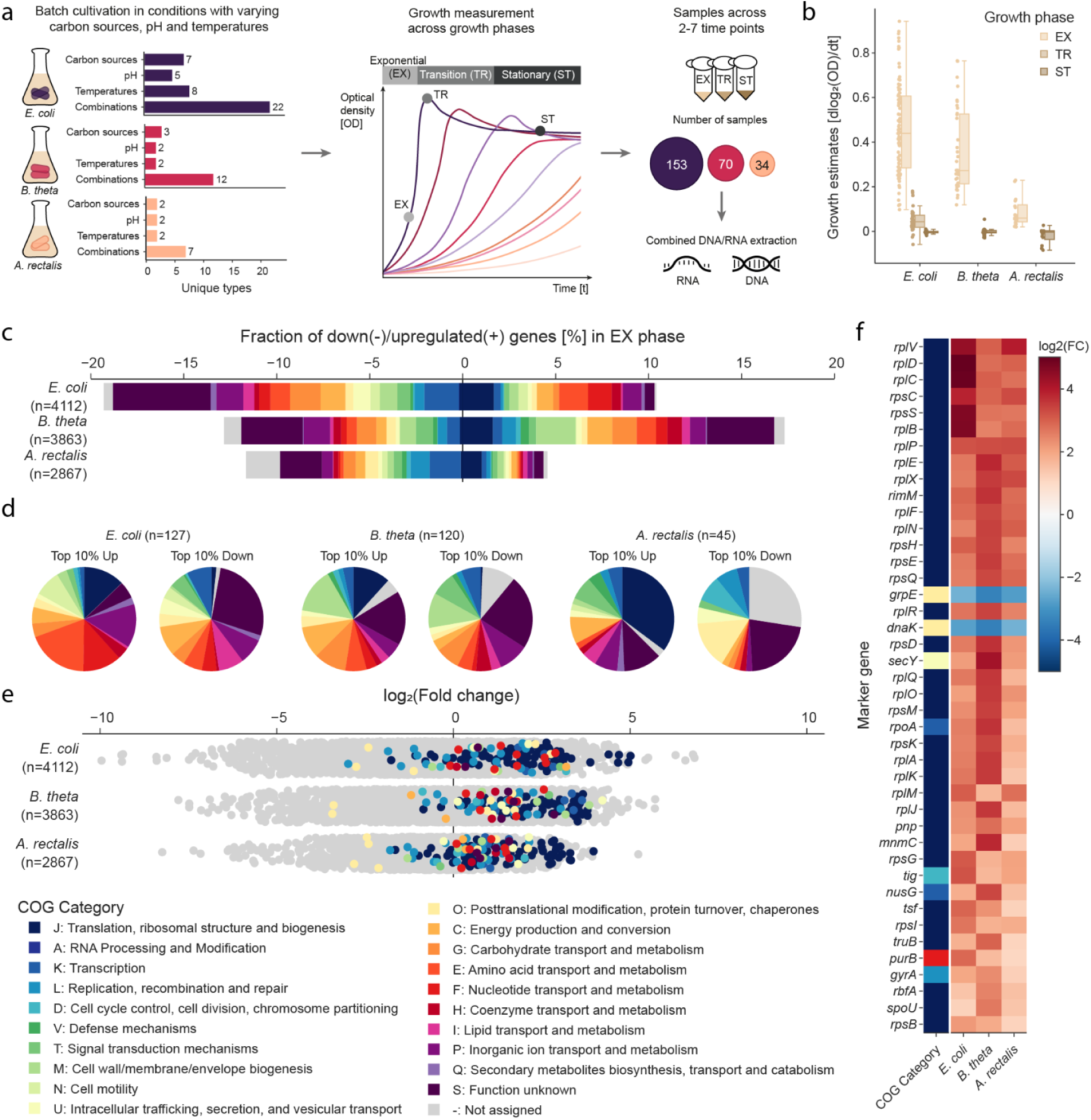
Growth phase-dependent transcriptomic signatures in bacterial strains from different phyla. a, Three bacterial strains were cultivated across combinations of diverse conditions. Growth was measured by changes in optical density (OD) over time (i.e., dlog_2_(OD)/dt) during exponential (EX; log-scaled linear increase in OD), stationary (ST; stable OD with net zero growth) and transition (TR; between EX and ST) phase. Cells were harvested at 2-7 time points across growth phases for co-extraction of DNA/RNA. **b,** Growth rates based on measured optical density over time (dlog_2_(OD)/dt) were obtained across the different bacterial strains, cultivation conditions and growth phases. **c**, The fractions [%] of differentially expressed (DE) genes (padj < 0.05, abs(log_2_FC) > 2, either downregulated (-, left) or upregulated (+, right)) were determined in exponential (EX) compared to stationary (ST) growth phase in each strain, based on the total number of genes (n) after pre-filtering for genes with ≥ 10 normalized counts averaged across samples (i.e., baseMean ≥ 10 filtering). Upregulated and downregulated genes were further subdivided according to clusters of orthologous genes (COG) categories. Multiple COG categories assigned to one gene were counted as separate assignments. **d,** In each strain, the proportions of COG categories were compared among the top 10% up-/ and downregulated DE genes (by log_2_FC; n = number of genes indicated on top). **e**, The distributions of log_2_FCs across growth phases (EX/ST) were compared between single-copy MGs (colored by COG category) and all other genes (light grey; n = number of genes per strain indicated on the left) with minimal expression (by ≥ 10 normalized counts averaged across samples, baseMean ≥ 10 filtering). **f,** The 42 MGs with congruent growth phase-dependent expression signatures (consistently padj < 0.05 and log_2_FC either all > 1 or < -1) across strains (sorted by absolute mean log_2_FC) were compared according to log_2_FCs (colored in the range [-5,5]) and assigned COG categories (colored box on the left).

To assess global growth phase-dependent transcriptome shifts, we filtered out genes with low counts across samples (< 10 averaged counts) and assessed the fraction of differentially expressed (DE) genes (padj < 0.05, absolute log_2_-fold changes (log_2_FCs) > 2) in each strain. Within up-/ or downregulated genes, we evaluated the enrichment of Clusters of Orthologous Genes (COG) categories by performing Fisher’s exact tests with Benjamini Hochberg correction for multiple testing (Fig. 1c; Supplementary Table 2). In *E. coli*, 10% of the genes were up-/ and 19% downregulated (29% DE) in EX phase, with categories J (Translation, ribosomal structure and biogenesis; 70/240 genes; padj = 8.14e-16), E (Amino acid transport and metabolism; 65/293; 6.02e-09), F (Nucleotide transport and metabolism; 53/243; 9.02e-08), and N (Cell motility; 21/107; 1.46e-02) significantly enriched in upregulated genes. In downregulated genes, COG category G was enriched (Carbohydrate transport and metabolism; 70/253; 9.92e-03). In *B. theta*, 17% of genes were up-/ and 13% downregulated (30% DE), with COG categories J (64/167; 5.56e-10), M (Cell wall/membrane/envelope biogenesis; 84/332; 7.05e-04) and C (Energy production and conversion; 53/192; 1.13e-03) enriched among upregulated genes. COG categories P (Inorganic ion transport and metabolism; 57/297; 2.30e-02), T (Signal transduction mechanisms; 37/180; 2.59e-02) and O (Posttranslational modification, protein turnover, chaperones; 20/88; 4.35e-02) were enriched in downregulated genes. In the slow-growing strain *A. rectalis*, 5% of genes were up-/ and 12% downregulated (17% DE), with COG category J (30/172; 1.37e-09) enriched in upregulated genes and K (Transcription; 46/245; 2.32e-03) as well as O (19/77; 4.63e-03) in downregulated genes. Substantial gene fractions also remained uncharacterized (S: Function unknown, -: Not assigned), constraining insights into functional enrichment among DE genes.

To assess whether these observations also hold true for the strongest differentially expressed genes, we extracted the top 10% up/- or downregulated genes (according to log_2_FCs) in each strain and repeated COG enrichment analysis (Fig. 1d; Supplementary Table 2). In *E. coli*, the same COG categories were enriched among the strongest upregulated genes in EX phase (E: 22/110; 1.37e-02, F: 16/79; 2.39e-02, J: 16/78; 2.39e-02 and N: 10/43; 4.01e-02). Among the strongest downregulated genes, COG category I (Lipid transport and metabolism; 10/29; 5.67e-03) was enriched. In *B. theta*, there was no enrichment of COG categories among the strongest DE genes (marginal significance for category J in upregulated genes; 15/70; 5.38e-02). In *A. rectalis*, COG category J was enriched (17/33; 3.43e-09) in genes upregulated in EX phase. In summary, genes involved in translation and ribosome biogenesis (J) were consistently overrepresented in all strains among genes upregulated in EX phase (both in all and the top 10% DE genes), in line with strong links between ribosome biosynthesis and growth^36,44^. Protein folding and chaperone genes (O) were overrepresented in genes upregulated in ST phase in *B. theta* and *A. rectalis* (only in all and not in the top 10% DE genes). These results suggest the strongest and most consistent transcriptional requirement for ribosome biogenesis genes in EX phase and potential secondary or more diverse requirements for protein folding and chaperone genes in ST phase.

With the aim to develop a transcriptomic growth predictor independent of the availability of reference genomes and universally applicable to bacteria, we assessed the growth phase-dependent expression of the 127 MGs leveraged by the taxonomy databases GTDB^45^ and proGenomes2^46^ (Methods; Supplementary Table 3). In each strain, we compared the distribution of log_2_FCs of these MGs to the overall spread of log_2_FCs across all genes. While all log_2_FCs were in the range from -10 to +7, log_2_FCs of MGs were distributed in a narrower range from -3.5 to +5. The majority of these 127 MGs were significantly upregulated (padj < 0.05; log_2_FC > 1) in EX phase (*E. coli*: 85/10 up/downregulated, *B. theta*: 94/3 up/down, *A. rectalis*: 57/10 up/down) (Fig. 1e), in line with many MGs being associated with translation and ribosome biogenesis. To determine which MGs displayed transcriptional shifts consistent across strains, we assessed the 42 out of the 127 MGs with congruent differential expression (padj < 0.05; either log_2_FCs all > 1 or all < -1) in all strains. We saw that 81% (34/42) were assigned to category J, mostly annotated as ribosomal proteins, and consistently upregulated across strains (Fig. 1f). Meanwhile, incongruent MGs were not significantly enriched for any functional category (Supplementary Fig. 1). In stationary phase, the MGs *dnaK* and *grpE,* assigned to category O and involved in protein folding and repair^47,48^ were consistently upregulated.

### Growth state classification based on transcriptomic MG profiles

To evaluate the use of MG transcripts for reference genome-independent growth prediction, we re-normalized the MG counts disregarding counts from all other genes. To perform gene length normalization, we divided raw read counts of each MG by its gene length (in kilobases), resulting in reads per kilobase (RPK). To normalize for sequencing depth within the MGs, we divided the RPK values by a scaling factor (i.e., the sum of RPK values across all MGs in each sample divided by one million), resulting in Transcripts Per Million MG counts (i.e., TPM_mg). Lastly, TPM_mg counts were log_2_-scaled after adding a pseudo-count of 0.5, resulting in transcriptomic MG profiles (Fig. 2a). First, we focused on the data generated for *E. coli*, as they encompassed the largest number of experimental conditions and samples (Fig. 1a). Applying a Principal Component Analysis (PCA) on the transcriptomic MG profiles of all *E. coli* samples, there was a clear separation of EX and ST samples along the first PC (PC1; explained variance = 59%) with TR samples distributed along a trajectory and partially overlapping with EX to ST samples (Fig. 2b). In addition to growth phase, differences in pH were driving a separation of samples along the second PC (Supplementary Fig. 2a) and, apart from PC3 (explained variance = 12%), the additional explained variance per subsequent PC (PC4 - PC20; explained variance each < 4%) was relatively low (Supplementary Fig. 2b).

**Figure 2.**
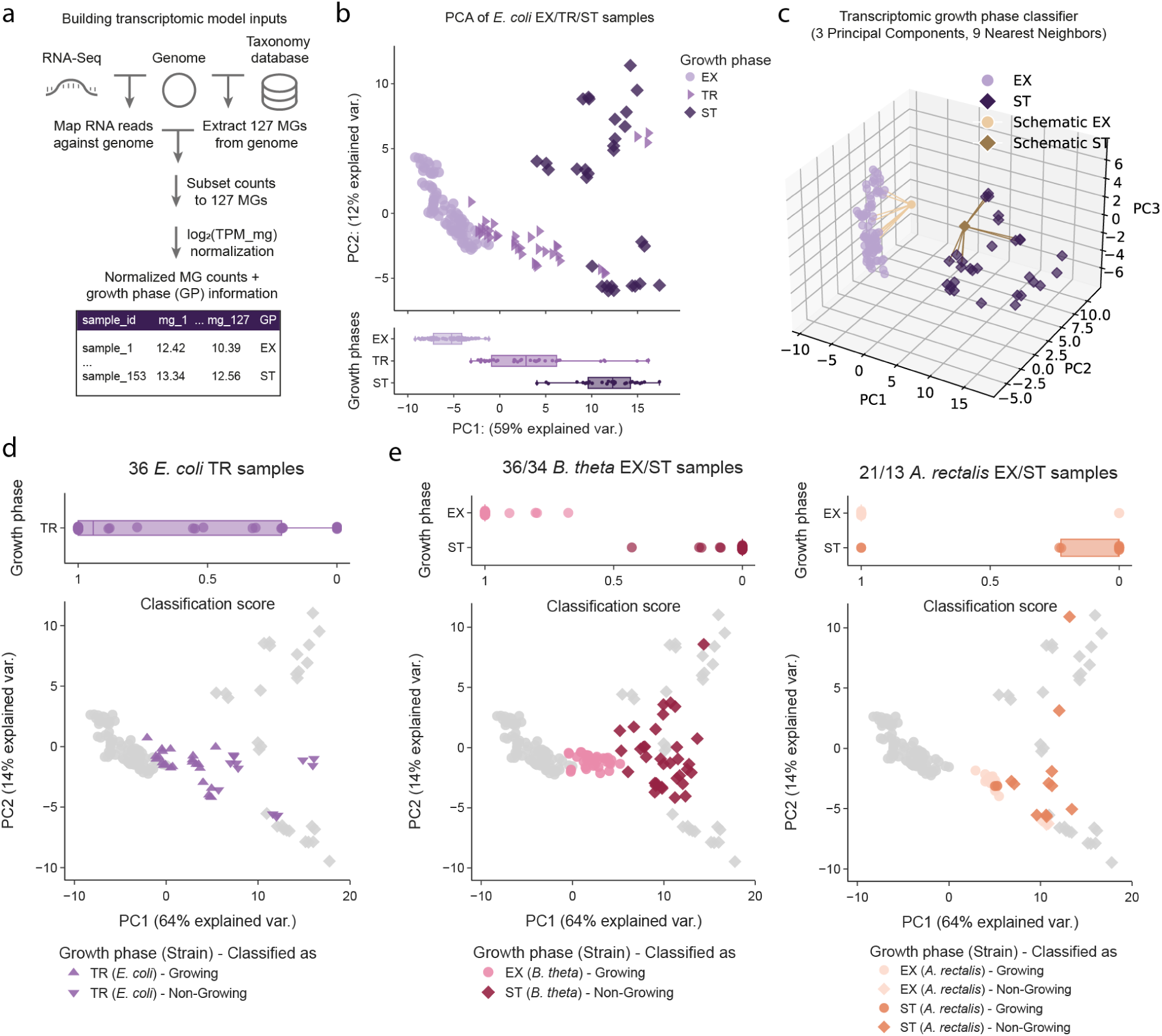
A transcriptomic growth classifier transferable across bacterial strains from different phyla. **a**, Raw RNA sequencing reads from *E. coli* samples were mapped against its reference genome. 129 MGs as defined by the GTDB^45^ (120) and proGenomes^46^ (40) databases were extracted from the reference genomes of the three strains using tools provided by database developers and subset to the 127 MGs shared across the three strains. The transcriptomic MG counts were normalized (log_2_(TPM_mg); Methods), yielding transcriptomic profiles with matching actual (i.e., experimentally determined) growth phases. **b,** A Principal Component Analysis (PCA) was performed on transcriptomic MG profiles of *E. coli* EX/TR/ST samples. The value ranges along PC1 are displayed as boxplots for each growth phase at the bottom. **c,** A binary growth phase classifier (3 PCs, 9 kNNs, metric=euclidean and weights=distance) was trained with transcriptomic MG profiles from *E. coli* EX/ST samples. **d**, Transcriptomic MG profiles of *E. coli* TR samples were transformed using three PCs derived from *E. coli* EX/ST data and classified as growing/non-growing using the PC, kNN-based classifier. Top panel: Boxplots depicting the distribution of classification scores obtained from the model, on an inverted x-axis ranging from 1 (growing) to 0 (non-growing). Bottom panel: Mapping of TR samples onto the first two PCs derived from *E. coli* EX/ST samples (light grey). Shapes correspond to classified growth states (Growing: triangle-up, Non-Growing: triangle-down). **e,** *B. theta* (left subpanel) and *A. rectalis* (right subpanel) samples were transformed using three PCs derived from *E. coli* EX/ST data and classified as growing/non-growing using the PC, kNN-based classifier. Top panel: Boxplots depicting the distribution of classification scores obtained from the model, on an inverted x-axis ranging from 1 (growing) to 0 (non-growing). Bottom panel: Mapping of *B. theta* and *A. rectalis* samples onto the first two PCs derived from *E. coli* EX/ST samples (light grey). Colors correspond to experimentally assigned growth phases (*B. theta*: red, *A. rectalis*: orange) and shapes correspond to classified growth states (Growing: circle, Non-Growing: diamond).

Given the separation of EX/ST samples according to transcriptomic MG profiles, we tested the predictability of growth (i.e., growing vs non-growing) by a non-parametric, supervised machine learning approach (i.e., k-Nearest Neighbor (kNN) model on PCs) (Fig. 2c). To this end, we trained a binary classifier (3 PCs, 9 kNN) on *E. coli* samples from EX and ST growth phase (EX: 87, ST: 30) with 5x cross validation (training/test data = 80/20%). The classifier returns scores from 0 to 1 and classifies bacteria as non-growing (scores < 0.5) or growing (scores >= 0.5) with a default threshold at 0.5. To assess how shifts in transcriptomic profiles during transition (TR) from EX to ST growth phase affect binary growth classification, we transformed the transcriptomic MG profiles of *E. coli* TR samples using the principal components derived from EX/ST samples and applied the classifier. We observed that TR samples obtained more widely distributed classification scores in the range from 0 to 1 (Fig. 2d). We thus interpreted classification scores as probabilities of growth ranging from 0 (non-growing) to 1 (growing) with values in between indicating larger uncertainty of the predicted growth state.

To test the generalizability of transcriptomic growth prediction, we applied the binary classifier to EX/ST samples collected for the other two strains (*B. theta* and *A. rectalis*). When transposed and embedded along the first two PCs derived from *E. coli* EX/ST data, *B. theta* and *A. rectalis* samples yielded a growth phase separation similar to *E. coli* (Fig. 2e). The majority of classification scores obtained for EX/ST samples were exactly 1 or 0 in *B. theta* (32/36, 29/34) and *A. rectalis* (18/21, 9/13). Despite some intermediate scores in *B. theta*, all EX/ST samples were classified correctly. In *A. rectalis*, five samples were misclassified (i.e., 3 EX samples classified as non-growing, 2 ST samples classified as growing). To explore the reasons for misclassification, we performed hierarchical clustering on whole transcriptomes and found that these samples clustered closest with transcriptomes from the classified growth states instead of from the experimentally assigned growth phases (Methods; Supplementary Fig. 3). Since the slow-growing strain *A. rectalis* displayed more heterogeneity in growth curves and between replicates, the OD-based growth phases were likely experimentally misassigned. Nevertheless, the classifier correctly classified all *B. theta* and *A. rectalis* samples which had conclusive experimental growth phase assignments, suggesting that the transcriptomic MG profiles contained growth phase-dependent signatures congruent across strains from different phyla. These results imply transferability of transcriptomic, MG-based growth phase prediction across the strains tested in this study.

### Comparison to PTR-based growth prediction

To assess how genomic PTR-based methods predict growth in our *in vitro* data across bacterial strains, we computed log_2_(PTR) values from genomic sequencing coverage for the same EX/ST samples using CoPTR^29^. As PTR-based tools were developed to predict growth rates, we first assessed the correlations of log_2_(PTR) estimates to measured growth rates in EX samples. While at 37 °C, log_2_(PTR) estimates correlated to measured growth rates in all strains (*E. coli*: R^2^ = 0.86; *B. theta*: 0.91; *A. rectalis*: 0.74), at decreased cultivation temperatures (33-20 °C) log_2_(PTR) values overestimated measured growth rates in EX phase, both in *E. coli* and *B. theta* (Supplementary Fig. 4a). Residuals to a linear model fitted to *E. coli* samples at 37 °C increased up to 7-fold at decreased cultivation temperatures (25-20 °C) leading to a potential bias of up to 0.35 in genomic log_2_(PTR) estimates (Supplementary Fig. 4b). While no bias was apparent for slow growth rates ≤ 0.2 h^-1^ in all three strains, large temperature-dependent log_2_(PTR) biases were observed for faster growth rates.

Next, we compared log_2_(PTR) value ranges across growth phases. In EX samples, the ranges of log_2_(PTR) values (*E. coli*: 0.42-0.94, *B. theta*: 0.33-1.12, *A. rectalis*: 0.24-0.59) were higher than growth rates measured by dlog_2_(OD)/dt in all strains (*E. coli*: 0.10-0.94 h^-1^, *B. theta*: 0.12-0.76 h^-1^, *A. rectalis*: 0.02-0.23 h^-1^) (Supplementary Fig. 4c). Despite the narrow ranges of OD-based growth rates close to zero in ST samples (-0.08-0.05 h^-1^), log_2_(PTR) values were distributed across considerably large log_2_(PTR) ranges (*E. coli*: 0.04-0.37, *B. theta*: 0.01-0.31, *A. rectalis*: 0.07-0.37) in ST samples, similar to ranges in EX samples. Consequently, log_2_(PTR) estimates would require calibrating and defining strain-specific thresholds for *E. coli* and *B. theta* to distinguish growing from non-growing cells. However, this would not be possible for the slow-growing strain *A. rectalis*, as log_2_(PTR) ranges of EX/ST samples were overlapping. Overall, these results indicate limited applicability of log_2_(PTR) estimates in predicting growth rates across temperature gradients and in distinguishing growth phases of slow-growing bacteria. Our tool therefore offers a complementary approach to assessing microbial community dynamics.

### Growth state associations of individual MGs in E. coli and across strains

Having trained a classifier on PCs of transcriptomic MG profiles of *E. coli*, we sought to evaluate which individual MGs drive the separation along PCs and therefore are likely associated with the different growth states (i.e., growing/non-growing). As there was a clear separation between growth phases along PC1, we identified the top 15 MGs with the largest absolute PC1 loadings in *E. coli* for closer inspection. Within these MGs, those associated with growing *E. coli* (Fig. 3a; negative PC1 loadings) were all ribosomal protein genes (*rplD*, *rplC*, *rplB*, *rpsS*, *rplV*) encoding for the proteins L4, L3, L2, S19 and L22. This finding is in line with previously identified proteins essential for ribosome assembly^49,50^. However, the ribosomal protein gene *rplT* encoding bL20 was associated with non-growing *E. coli* (Fig. 3a; positive PC1 loadings) in line with it being observed as downregulated in EX phase according to differential expression (DE) analysis (Supplementary Fig. 1). Ribosomal protein bL20 is also among the proteins essential for the first *in vitro* reconstitution step of the large ribosomal subunit but was shown to be non-essential for ribosome activity as it can be withdrawn from the mature 50S subunit^51^. Furthermore, bL20 has been described to repress the translation of its own operon *infC-rpmI-rplT* by binding to its mRNA, suggesting it to be an autogenous repressor in *E. coli*^52–55^ and *B. subtilis*^56^. Other MGs associated with non-growing *E. coli* (Fig. 3a; positive PC1 loadings) encode proteins involved in protein folding (e.g., *dnaK*, *grpE*), proteolysis cascades (e.g., *clpX*), DNA replication and repair (e.g., dnaG, *recN*, *uvrB*, *ruvA*/*ruvB*) and ribosomal RNA methylation (e.g., *rsmH*). The respective proteins carry out important functions in handling denatured proteins^47^ or DNA damage^57–59^ during stress/SOS response induced by e.g., hyperosmotic/heat shock or UV light. Similarly, entry into stationary phase and associated nutrient deprivation in batch cultures increase levels of denatured proteins, oxidative stress and DNA damage and trigger RpoS-mediated stress response^60^. For some of these MGs, their direct relevance during ST phase has previously been reported in *E. coli* (e.g., *dnaK*^48^, *clpX*^61^).

**Figure 3.**
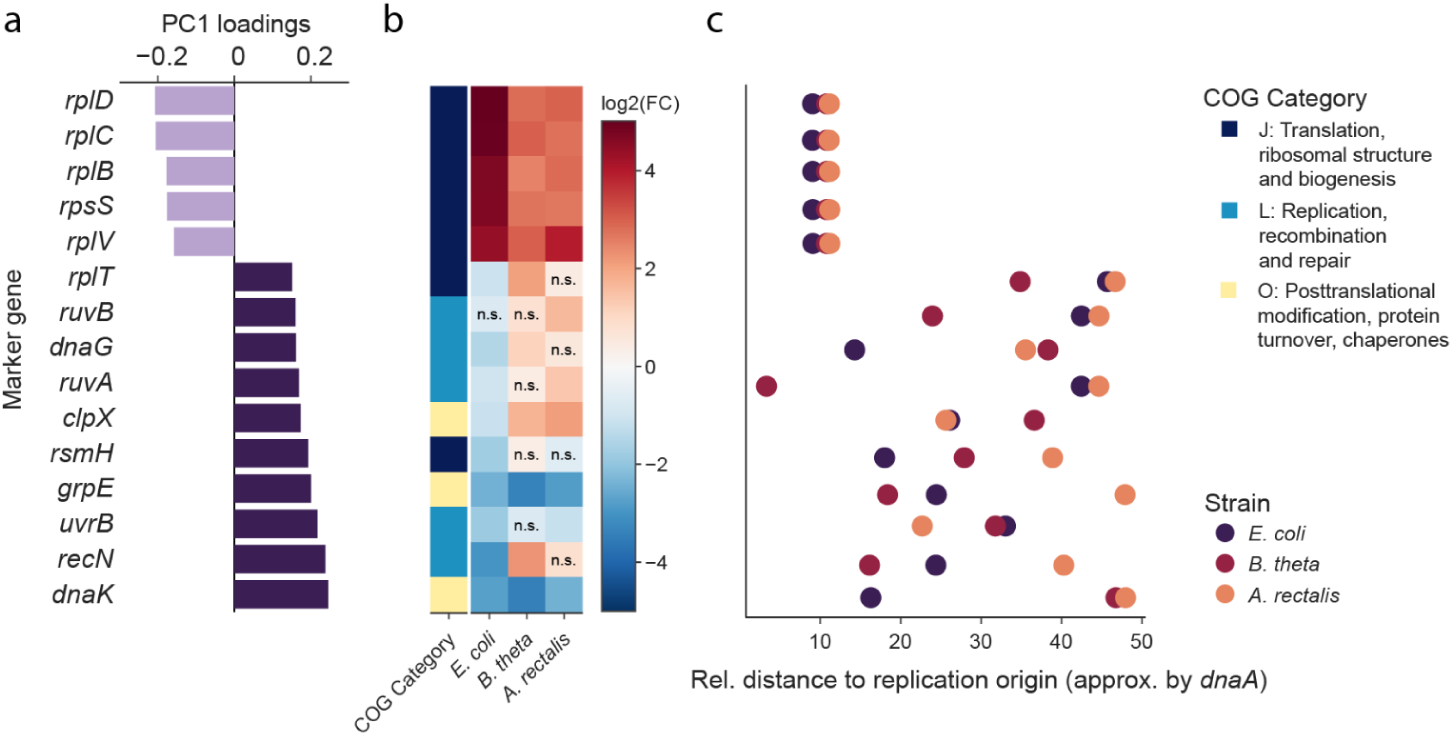
The MGs with largest absolute PC1 loadings in *E. coli* with respect to their COG categories, growth phase-dependent expression levels and genomic location. a, The PC1 loadings for the 15 MGs with largest absolute PC1 loadings were derived from the PCA on transcriptomic MG profiles of *E. coli* EX/ST samples. **b**, The log_2_FCs across EX/ST growth phases based on differential expression analysis with PyDESeq2^63^ were compared between strains for each of the 15 MGs with largest absolute PC1 loadings in *E. coli* (n.s. = non-significant, by adjusted p-value using Benjamini-Hochberg correction >= 0.05). The assigned COG category is depicted in the leftmost column (J: dark blue, L: light blue, O: yellow). **c,** Relative distance to replication origin of these 15 MGs (approximated by *dnaA* at 0) within each circularized reference genome (*E. coli*: purple, *B. theta*: red, *A. rectalis*: orange) were compared between strains.

As the classifier correctly predicted the growth states in the other two strains, we wanted to test whether the growth phase associations of these 15 MGs in *E. coli* would readily apply to the other two strains (*B. theta* and *A. rectalis*). To this end, we compared log_2_FCs across growth phases according to DE analysis between the strains (Fig. 3b). Most ribosomal protein genes (*rplD*, *rplC*, *rplB*, *rpsS*, *rplV*) belonging to COG category J (Translation, ribosomal structure and biogenesis) and two protein/DNA repair genes (*dnaK* and *grpE*) belonging to COG category O (Posttranslational modification, protein turnover, chaperones) were differentially expressed (padj < 0.05, abs(log_2_FC) >= 1) with congruent directionality across strains. Meanwhile, other MGs (*recN*, *uvrB*, *rsmH*, *clpX*, *ruvA*/*ruvB*, *dnaG* and *rplT*) with a majority belonging to COG category L (Recombination, replication and repair) displayed opposing strain-specific log_2_FCs or non-significant differences between growth phases in at least one strain (Fig. 3b). These findings suggest that some MGs identified as growth state-dependent in *E. coli* may have limited generalizability in differential expression across bacterial strains from different phyla, or may be affected by the proportional nature of transcriptomic profiles. This observation emphasizes the importance of building a classifier which considers multivariate signatures and the relationships between MGs.

To determine whether growth state-associated MGs show conserved positional bias across bacterial strains, we assessed whether the relative distance to the origin of replication (*ori*; in % of the whole circularized genome with *ori* approximated by the location of *dnaA*) of these 15 MGs was similar across strains. Ribosomal protein genes associated with growing states were located relatively close to the replication origin (as expected according to replication-based gene dosage effects resulting in biased localization of genes involved in transcription and translation towards the *ori*^62^). Meanwhile, MGs associated with non-growing states were more widely distributed in distance to the replication origin, and the genomic position was not conserved between strains. (Fig. 3c). These results suggest that genomic location alone may be unsuitable to identify MGs associated with non-growing states and that a data-driven approach to identify growth phase-associated MGs may be superior to relying on genes in pre-determined functional groups or within specific genomic locations.

### In vivo classification of growth states in EAM mice

To test the applicability of our developed binary growth classifier in a community context *in vivo*, we explored the diurnal dynamics in gnotobiotic Easily Accessible Microbiota (EAM) mice containing the strains studied here *in vitro* (*E. coli*, *B. theta* and *A. rectalis*). We collected cecum content samples across three daytimes (2, 5 and 9 pm) from four mice per daytime and extracted DNA and RNA for sequencing (Fig. 4a; Supplementary Table 4; one 2 pm sample was disregarded due to insufficient biosample and data quality). These samples span the shift from day to night, during which the food intake of the mice increases, as they are nocturnal animals. The increased food available to the gut microbiota may lead to higher microbial metabolic activity and growth, coinciding with rising hydrogen measurements in the exhalome^64^. To assess potential shifts in community composition across daytimes, we quantified absolute cell numbers (per gram of dry weight; Methods) by qPCR. *B. theta* and *A. rectalis* were present at similar numbers (4.5*10^11^ cells/g dry weight), whereas *E. coli* was present at lower numbers (1*10^11^ cells/g dry weight). Based on these cell numbers, we inferred relative abundances (*E. coli*: 10%, *B. theta*/*A. rectalis*: 45%) across samples and daytimes (Fig. 4b) implying stable community dynamics.

**Figure 4.**
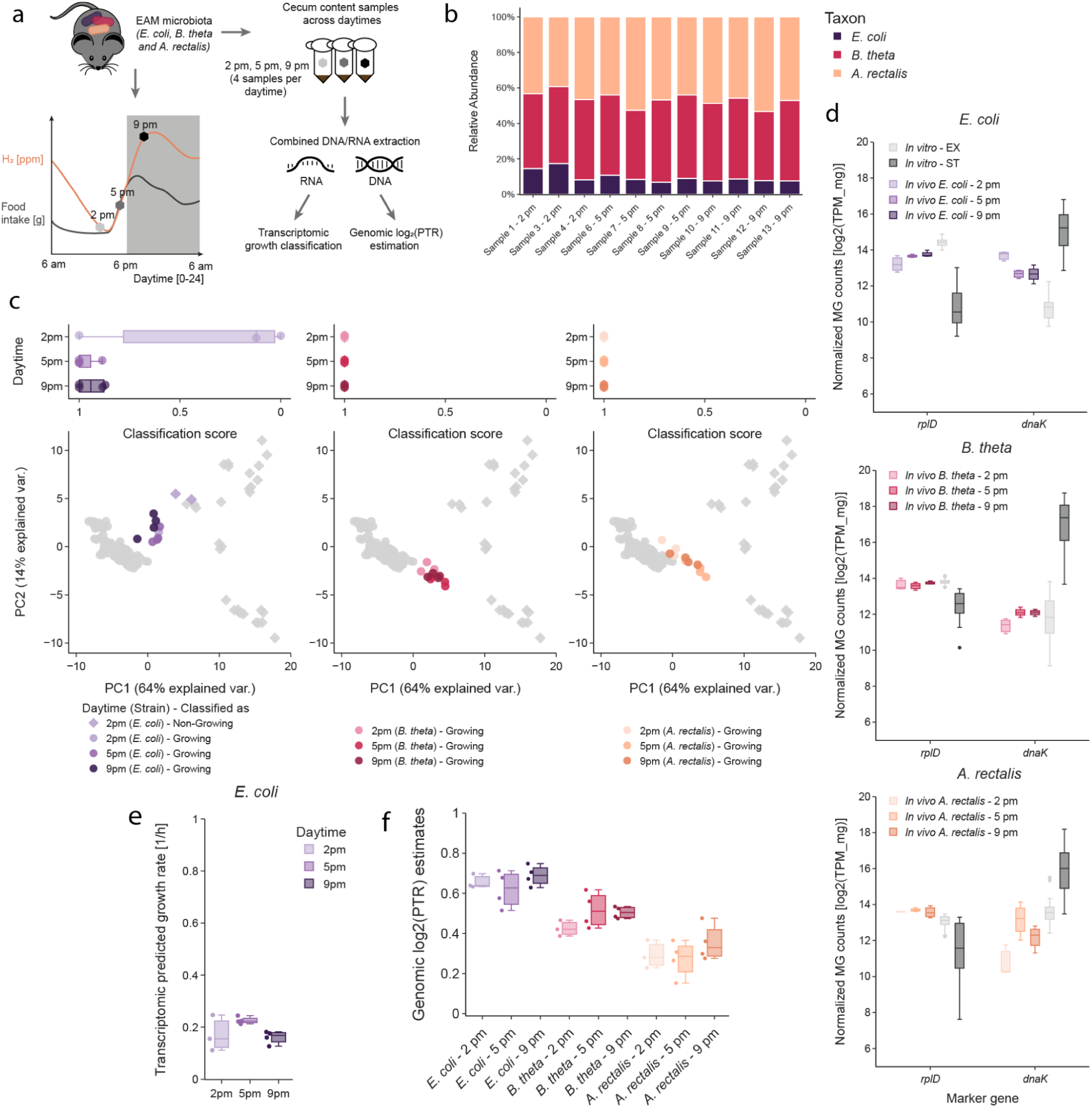
*In vivo* metatranscriptomic, marker gene-based growth prediction in EAM mice. a, We generated paired metagenomic/metatranscriptomic data from cecum content of EAM mice (with a gut microbiota consisting of *E. coli*, *B. theta* and *A. rectalis*) across three daytimes (2 pm, 5 pm, 9 pm) coinciding with food intake and H_2_ measurements in the exhalome. **b,** We inferred relative abundances from absolute cell numbers of the EAM strains per gram of dry weight of cecum content by qPCR using strain-specific primers and standard curves. **c,** *In vivo* transcriptomic MG profiles of the EAM strains were transformed using three PCs derived from *E. coli in vitro* EX/ST data and classified as growing/non-growing using the PC, kNN-based classifier. Top panel: Boxplots depicting the distribution of classification scores obtained from the model, on an inverted x-axis ranging from 1 (growing) to 0 (non-growing). Bottom panel: Mapping of *E. coli*, *B. theta* and *A. rectalis* transcriptomic MG profiles from cecum content samples onto the first two PCs derived from *E. coli in vitro* EX/ST samples (light grey). Colors correspond to the EAM strains (*E. coli*: purple, *B. theta*: red, *A. rectalis*: orange) and shades correspond to the daytimes of collected cecum content (light: 2 pm, medium: 5 pm, dark: 9 pm). Shapes correspond to classified growth states (Growing: circle, Non-Growing: diamond). **d,** The normalized transcriptomic counts (log_2_(TPM_mg)) of the two MGs with largest PC1 loadings in *E. coli* (*rplD*, *dnaK*) were compared between *in vivo* and *in vitro* EX/ST samples. **e,** *In vivo E. coli* growth rates (h ^-1^) were predicted based on a transcriptomic Ridge regression model trained on *in vitro E. coli* EX/TR/ST samples. **f,** Genomic-based log_2_(PTR) estimates for each strain across daytimes.

To predict *in vivo* growth states, we generated metatranscriptomic data, computed transcriptomic MG profiles in each strain and applied our growth classifier (Fig. 4c). *E. coli* received intermediate classification scores ranging from 0 to 1, and was classified as non-growing in 2/3 samples at 2 pm and as growing in all other samples. The other two strains (*B. theta*, *A. rectalis*) were classified as growing with classification scores being exactly 1 in all samples. To evaluate whether *in vivo* transcriptomic MG profiles of the two MGs with strongest PC1 loadings in *E. coli* (*rplD*, *dnaK*) recapitulated *in vitro* profiles, we compared their normalized transcriptomic counts (log_2_(TPM_mg); Methods) between *in vivo* and *in vitro* samples in each strain (Fig. 4d). In *E. coli*, *in vivo* counts were distributed between EX and ST *in vitro* counts in line with the obtained intermediate growth classifications. In contrast, *in vivo* counts in *B. theta* and *A. rectalis* aligned with EX *in vitro* counts, supporting their classifications as growing across all daytimes. In *A. rectalis*, the *in vivo* counts of *dnaK* were lower than in *in vitro* EX samples. This may be explained by *A. rectalis* growing slowly in minimal media, whereas it is a dominant community member in EAM mice (together with *B. theta*)^64^ and likely growing at higher rates *in vivo*.

To further support the obtained growth classifications, we performed hierarchical clustering of *in vivo* and *in vitro* whole transcriptomes for each strain (Supplementary Fig. 5a). Indeed, *in vivo* whole transcriptomes of *B. theta* and *A. rectalis* clustered with *in vitro* EX transcriptomes. In contrast, those of *E. coli* clustered closest with *in vitro* TR transcriptomes across all daytimes, suggesting that *E. coli* in the mouse gut may be in a physiological state dissimilar from the EX or ST phases *in vitro*. To further investigate the lower classification confidence for *E. coli*, we leveraged the large sample number and wide range of measured rates (-0.06 - 0.94 h^-1^) of *E. coli* across all growth phases (EX/TR/ST) with the aim to go beyond classifying growth states and predict actual rates. We trained multiple regression models (Methods; Supplementary Table 5) with 5x cross validation (training/test data = 80/20%) on all *E. coli* samples (EX = 87, TR = 36, ST = 30) and evaluated model performance (i.e., by mean absolute error (MAE) and R-squared) (Supplementary Fig. 5b). While all models displayed similar performance, the Ridge regressor performed best (MAE: 0.07, R-squared: 0.88). We thus re-trained the Ridge regression model on *E. coli* samples from all growth phases (EX/TR/ST) and applied it to predict growth rates for *E. coli in vivo* (Fig. 4e), resulting in relatively slow predicted growth rates (0.11 - 0.25 h^-1^) in all samples. These results suggest that our transcriptomic, marker gene-based growth classifier is transferable from *in vitro* isolate cultures to *in vivo* community settings in the EAM microbiota. While *B. theta* and *A. rectalis* were predicted as consistently growing, *E. coli* may undergo distinct or more complex growth dynamics, reflected in higher uncertainty of the classifier.

To evaluate existing PTR-based methods *in vivo*, we generated metagenomic sequencing data from the same samples and computed log_2_(PTR) estimates (Fig. 4f). While we also observed no significant changes of log_2_(PTR) estimates across daytimes, we obtained strain-specific log_2_(PTR) ranges with the highest range for *E. coli* (0.51-0.75). PTR-based growth rate estimates obtained for *E. coli in vivo* were within the range of EX samples *in vitro* (0.42-0.94). However, log_2_(PTR) estimates of *in vitro* TR samples (0.36-0.52 at pH 7.0 representative of the EAM cecum^64^; 0.11-0.40 at pH < 7.0) and ST samples (0.04-0.37) were substantially higher than expected based on the observed slow OD-based growth rates (TR: -0.05-0.18 h^-1^, ST: -0.02-0.01 h^-1^; Supplementary Fig. 5c), These results showcase the challenges in distinguishing growth phase shifts according to log_2_(PTR) estimates *in vitro* and *in vivo*. These results suggest that assessing growth *in vivo* with more factors at play and more diverse physiological states may be more challenging to assess with both metagenomic and metatranscriptomic growth prediction. However, our classifier may increase interpretability by providing classification scores that enable evaluating model uncertainty.

### Assessment of lipopolysaccharide-associated growth inhibition in Oligo-MM^12^ mice

To evaluate transcriptomic growth state classification beyond the three strains tested *in vitro*, we applied the classifier to strains from three other species within gnotobiotic mice containing a 12-member mouse gut community (Oligo-MM^12^). In a previous study^43^, sub-lethal intravenous lipopolysaccharide (LPS) or phosphate buffer saline (PBS) control injections were administered to six mice each to assess microbiota perturbations triggered by systemic inflammatory response. Perturbation effects on growth were assessed within metatranscriptomic data from cecum content six hours post injection (Fig. 5a; data obtained from <u>PRJEB82444</u>). In the respective study, LPS treatment led to strong multifactorial microbiota perturbations and differential expression analysis performed for three strains (*Enterocloster clostridioformis* YL32, *Blautia pseudococcoides* YL58, and *Bacteroides caecimuris* I48) revealed downregulation of amino acid biosynthesis genes and upregulation of oxidative stress and protein folding genes in all strains, as well as downregulation of ribosome biosynthesis genes in YL32 and YL58. Thereby, the conducted study^43^ suggests activation of stress response and potential LPS-associated growth inhibition in some microbial community members.

**Figure 5.**
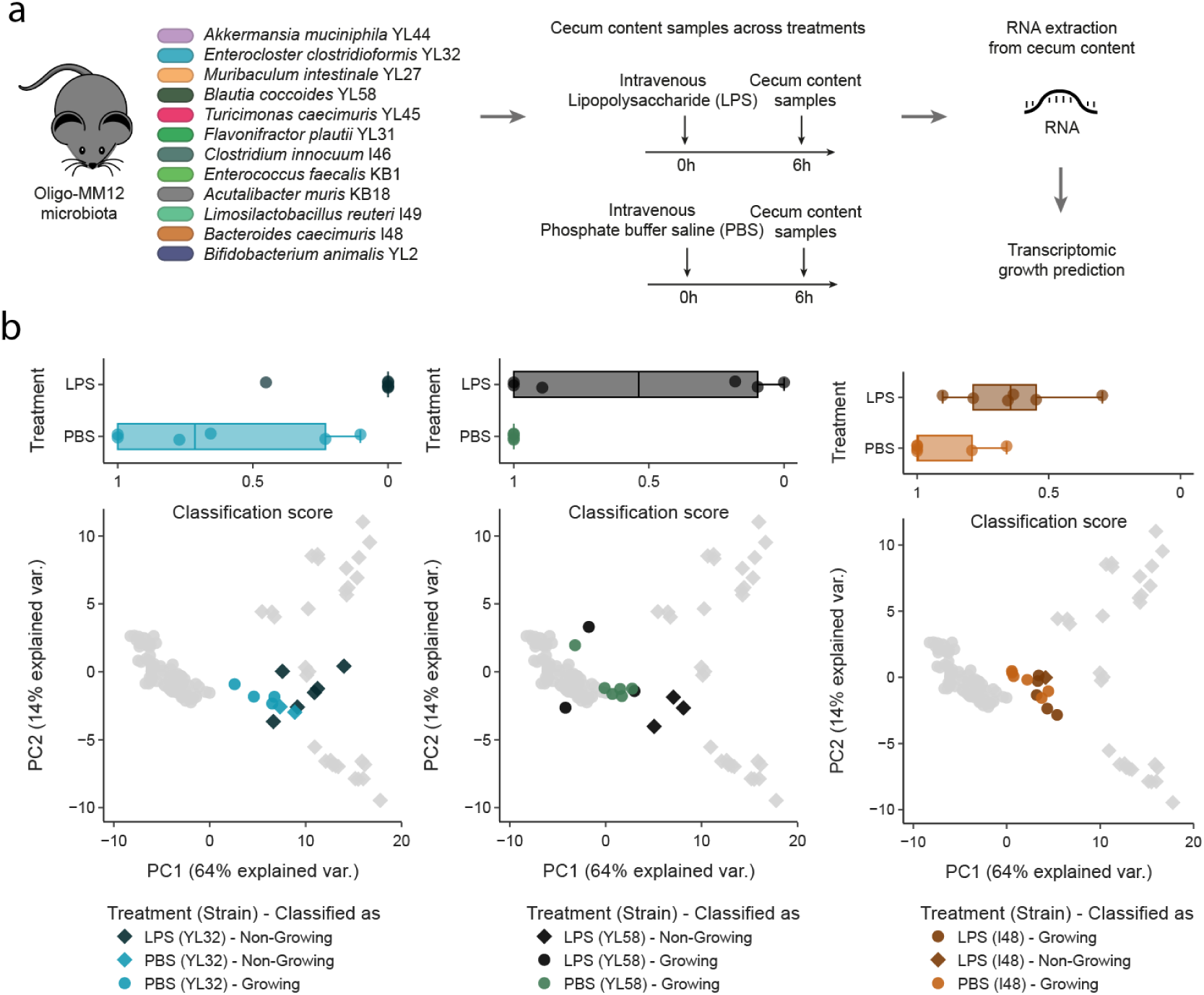
*In vivo* metatranscriptomic growth prediction to study growth dynamics upon lipopolysaccharide treatment in Oligo-MM^12^ mice. a, The previously conducted study^43^ treated Oligo-MM12 mice with sub-lethal intravenous injection of lipopolysaccharide (LPS) or phosphate buffered saline (PBS) control and generated metatranscriptomes from cecum content 6 hours post treatment. b, *In vivo* transcriptomic MG profiles of the three Oligo-MM^12^ strains of interest (*Enterocloster clostridioformis* YL32, *Blautia pseudococcoides* YL58 and *Bacteroides caecimuris* I48) were transformed using three PCs derived from *E. coli in vitro* EX/ST data and classified as growing/non-growing using the PC, kNN-based classifier. Top panel: Boxplots depicting the distribution of classification scores obtained from the model, on an inverted x-axis ranging from 1 (growing) to 0 (non-growing). Bottom panel: Mapping of YL32, YL58 and I48 transcriptomic MG profiles from cecum content samples onto the first two PCs derived from *E. coli in vitro* EX/ST samples (light grey). Colors correspond to the three strains (YL32: turquoise, YL58: olive green, I48: brown) and shades correspond to the treatment (dark: lipopolysaccharide (LPS) treatment, light: PBS control). Shapes correspond to classified growth states (Growing: circle, Non-Growing: diamond).

In agreement with the suggested growth inhibition, these strains obtained lower classification scores in cecum samples from LPS-treated mice compared to PBS controls (Mann-Whitney U test without correction for multiple testing; YL32: p-value = 0.009, YL58: 0.028, I48: 0.032) (Fig. 5b). YL32 was predicted as non-growing in 6/6 LPS-treated samples and as growing in 4/6 PBS controls. YL58 was predicted as non-growing in 3/6 LPS-treated samples and as growing in 6/6 PBS controls. I48 was predicted as non-growing in only 1/6 LPS-treated sample and as growing in 6/6 PBS controls. Notably, obtained classification scores were more widely distributed in the range from 0 to 1, representing larger classification uncertainty, which could originate from variability in treatment effects between mice (e.g., considerable mouse-to-mouse variation in triggered immune responses), more diverse physiological states or could indicate a requirement for more training data. Nevertheless, the growth classifications (into growing/non-growing) reproduced previously suggested growth inhibition according to differential expression analysis in these three strains.

## Discussion

In this study, we assessed transcriptomic signatures of bacteria cultured *in vitro* and evaluated the use of 127 MGs for predicting growth states according to growth phases. Trained on *E. coli* transcriptomic MG profiles originating from diverse cultivation conditions, our classifier was able to predict the growth states of *B. theta* and *A. rectalis* under varying carbon sources, pH, and temperature. This suggests that transcriptomic, MG-based growth prediction has the potential to generalize across taxa under diverse environmental conditions. Furthermore, we demonstrated how our method complements metagenomic log_2_(PTR)-based growth prediction, which would have required determining strain-specific thresholds to distinguish growth phases in the two fast-growing strains (*E. coli*, *B. theta*). For the slow-growing strain *A. rectalis*, log_2_(PTR)-based growth phase distinction was not possible at all. These findings have important implications, as the majority of microbial community members may be non-growing or slow-growing for the majority of time depending on the environment^35,65^ and obtained log_2_(PTR) estimates may misrepresent slow-growing community members in a non-growing state as growing. Moreover, log_2_(PTR) estimates, when used to predict actual growth rates, overestimated growth rates in decreased cultivation temperatures in *E. coli* and *B. theta*. This observation may arise from, for example, decreased enzymatic activity of the replication machinery at lower temperatures required for balanced coordination of cell division and DNA replication^66^, which may render replication-based signatures unsuitable to predict bacterial growth rates across temperature gradients that are central to certain environments (e.g. soil, aquatic systems).

Upon investigating transcriptomic signatures of individual marker genes, we found ribosomal protein genes (e.g., *rplD*, *rplC*, *rplB*, *rpsS*, *rplV*) and DNA/protein repair genes (e.g., *dnaK, grpE*) to be among the MGs with largest PC1 loadings in *E. coli* and with the most congruent differential expression across strains. These MGs are likely important features in the classifier as their expression is associated with growth phases in *E. coli*, consistent with prior studies^36,48^. Moreover, these growth-state related expression patterns appear to generalize across strains, as many of these MGs showed consistent differential expression patterns in *B. theta* and *A.rectalis*. However, we observed limited generalizability of growth phase-dependent expression for some MGs. Notably, some ribosomal protein genes (e.g. *rplT, rplU*) were incongruent across strains and not significantly different in expression across growth phases. At the same time, MGs belonging to other functional categories (e.g. *dnaK*, *grpE*) were congruent in their differential expression across strains and informative of non-growing states. Nevertheless, our binary growth classifier was sufficiently robust to correctly classify all samples across strains. While biases in genomic location have been suggested to be present for many functional groups of genes^67^, we observed that particularly MGs informative of non-growing states had wider distributions of genomic locations, suggesting less constraints in chromosomal position from the origin of replication to the terminus. These results showcase how the data-driven development of a growth classifier may provide more accurate results than relying on genes in pre-determined functional groups or within specific genomic locations.

When applying the classifier *in vivo* in EAM mice, all strains, except for *E. coli* in two cecum samples, were predicted as growing in line with expected growth requirements of bacteria in the gut to prevent being washed out^68^. For *E. coli*, the wider distribution of classification scores and the clustering of whole transcriptomes with *in vitro* TR samples suggest that *E. coli in vivo* may experience distinct or more complex growth dynamics in the mouse gut. Some potential caveats of these results are that *E. coli* was cultivated aerobically *in vitro* (in contrast to the other two strains), which may result in the anaerobic conditions *in vivo* affecting transcriptomic MG profiles and growth classifications. Furthermore, *E. coli* was predicted as non-growing in the only two male mice in the study, suggesting potential gender effects. In contrast, *in vivo* metagenomic log_2_(PTR) estimates suggested high growth rates for *E. coli in vivo*, contradicting metatranscriptomic growth rate regression in *E. coli*. Lastly, by applying our classifier to a 12-member synthetic mouse gut community, we recapitulated previously suggested growth inhibition in three strains triggered by systemic inflammatory response^43^. The more widely distributed classification scores may indicate mouse-to-mouse variation in microbiota perturbations or limited generalizability of the model beyond the EAM strains. As a result, capturing additional physiological states and diverse biological contexts would likely further improve model performance.

In conclusion, this work advances bacterial growth prediction by developing a growth state classifier based on transcriptomic signatures of single-copy marker genes and testing transferability in three bacterial strains from different phyla. Previously, only one study predicted growth rates from whole transcriptomes in *E. coli*^38^, however without the potential for transferability across species. This study describes the first attempt to predict the growth states of diverse bacterial strains from different phyla in a wide range of comparable cultivation conditions and demonstrates applicability in microbial communities, thereby providing a novel method for *in situ* growth classification. Compared to existing methods for *in situ* bacterial growth prediction, our classifier displays i) broader applicability by mapping reads to marker genes rather than complete reference genomes and containing low dependence on assumptions due to data-driven modeling, ii) increased interpretability by providing classification scores and iii) improved accuracy by distinguishing growth states for slow-growing bacteria. Our results revealed that future work could focus on identifying a reduced set of MGs with congruent growth state-dependent transcriptomic signatures across strains to converge on a set of universal growth state-predicting marker genes. Furthermore, using more variable training data (e.g., expanding cultivation conditions, more diverse growth states, additional strains, diverse *in vivo* conditions) and/or exploring other data normalization approaches would likely improve classification accuracy and pave the way towards other types of classification (e.g., multiclass classifier to predict transitions between growth states). Nevertheless, our work demonstrates the potential of the method to generalize across phylogenetically diverse taxa, scale to more complex communities and be transferable to other environments. In combination with the information-rich nature of metatranscriptomic data, the classifier is envisioned to be informative to contribute to a more integrated understanding of microbial community dynamics and their effects on ecosystem processes upon environmental change.

## Methods

### In vitro experimental design and sample processing

#### Bacterial strains, liquid media and precultures

The bacterial strains *Escherichia coli* HS (CP092639.1), *Bacteroides thetaiotaomicron* VPI5482 (CP092641.1) and *Agathobacter rectalis* VPI0990 (CP092643.1) were revived from 50% cryostocks (50% Lysogeny Broth (LB) medium, 50% glycerol) stored at -80°C.

1. *E. coli* was streaked out on Lysogeny Broth (LB) agar and grown in the presence of oxygen overnight at 37 °C. For the preculture, a single colony was inoculated into 5 ml M9 minimal medium (12.8 g/L Na_2_HPO_4_·7H_2_O, 3 g/L KH_2_PO_4_, 0.5 g/L NaCl, 1 g/L NH_4_Cl, 100 μM CaCl_2_, 2 mM MgSO_4_) supplemented with Wolin’s trace elements (13.4 μM EDTA, 3.1 μM FeCl_3_-6H_2_O, 0.62 μM ZnCl_2_, 76 nM CuCl_2_-2H_2_O, 42 nM CoCl_2_-2H_2_O, 162 nM H_3_BO_3_, 8.1 nM MnCl_2_-4H_2_O), 20 mM Glucose and adjusted to pH 7.0. The media were sterilized with 0.22 μm filters. 16x160 mm culture tubes were used to incubate the aerobic precultures overnight at 37 °C while shaking at 180 rpm (Multitron, INFORS HT) and growing them to full density.
2. *B. theta* and *A. rectalis* were streaked out on Brain Heart Infusion (BHI) agar supplemented with 1 g/l Cysteine and 10% defibrinated sheep blood and were grown in the absence of oxygen overnight at 37 °C. For the preculture, single colonies were inoculated into 5 ml medium adapted^69^ from Tryptone Yeast Extract Glucose (TYG) medium (1% Tryptone, 100 mM K_2_HPO_4_/KH_2_PO_4_ (ratio adjusted for pH), 50 mM NaCl, 0.5mM CaCl_2_, 0.4 mM MgCl_2_, 50 µM MnCl_2_, 50 µM CoCl_2_, 4 µM FeSO_4_, 5 mM Cysteine, 20 mM NaHCO_3_, 5 mM Na_2_SO_4_, 20mM NH_4_Cl, 1.2 mg/l Hemin, 1mg/l Menadione, 2 mg/l Folinic acid and 2 mg/l Vitamin B12) and supplemented with 1 mg/l resazurin, 20 mM Glucose and adjusted to pH 7.0. The media were sterilized with 0.22 μm filters, introduced into the anaerobic tent at least 48h before inoculation to remove residual oxygen and covered with aluminium foil for light protection. 16x125 mm Hungate tubes were used to incubate the anaerobic precultures overnight at 37 °C while shaking at 180 rpm (Thermomixer, Eppendorf) and grow them to full density.

#### Batch culture conditions, growth measurement and harvesting

Across all cultivation experiments, the precultures were washed two times in 1x PBS, the optical density (OD) was estimated from OD measurement (OD600 reader, Labgene) of a 10-fold dilution in 1x PBS and the undiluted cells were added to each batch cultivation sample to achieve a starting OD of 0.03. A wide variety of sole carbon sources (Glucose, Maltose, Ribose, Arabinose, Fructose, Gluconate and Succinate), pH (5.5 - 7.0) and temperatures (20 - 37 °C) were tested (Supplementary Table 1; described in detail in following paragraphs). To this end, either 20 mM Glucose (C6) or other sole carbon sources were added at an equivalent carbon content to the base media. Varying pH in the media was achieved by addition of hydrochloric acid (HCl) or sodium hydroxide (NaOH). Per condition, a culture triplicate was grown per harvesting time point to ensure three biological replicates and an additional culture triplicate was monitored until the end of the experiment to obtain the full growth curve. The bacterial cells were harvested by centrifugation at 3500 x g for 3 min, supernatant removal and flash-freezing in liquid nitrogen. The culture pellets were stored at -80 °C until DNA/RNA extraction.

To investigate transcriptomic signatures across exponential growth rates, *E. coli* was cultivated in M9 minimal medium with varying carbon sources, pH (5.5 - 7.0), and temperatures (20 – 37 °C) as aerobic batch cultures. Per condition, 5 ml culture triplicates were incubated while shaking at 180 rpm (Multitron, INFORS HT) and OD measurements (OD600 reader, Labgene) were taken in 30 min (at >= 30°C) or 1h (at < 30°C) intervals from 50 µl subsamples added to 450 µl base medium. 1 ml of cell culture was harvested for each replicate per condition at approx. OD 0.5 during mid-exponential growth.

To investigate transcriptomic signatures across growth phases, *E. coli* was cultivated with a subset of conditions as aerobic 250 μl batch cultures. The culture triplicates were grown in 96 well format with fast 280 cpm shaking. A plate reader (BioTek Synergy H1 Multimode Reader, Agilent) was used for automated OD measurements in 10 min intervals. Per culture triplicate, the whole culture volume (250 μl) of each of the three samples was harvested at the respective timepoint (3-7 time points, including mid/late exponential (EX1/2), first plateau (PL), death (IH), slow growth (SG1/2) and stationary (ST1/2)).

To test the generalizability across other bacterial strains, *B. theta* and *A. rectalis* were cultivated in adapted TYG medium with different carbon sources (Glucose, Maltose), pH (5.9, 7.0), and temperatures (31, 37 °C) as anaerobic 250 μl batch cultures. The culture triplicates were inoculated using sterile syringes and needles, sealed with transparent membranes (BreatheEasy, Sigma Aldrich) and grown in 96 well format with fast 280 cpm shaking. A plate reader (BioTek Synergy H1 Multimode Reader, Agilent) was used for automated OD measurements in 10 min intervals. Per culture triplicate, the whole culture volume (250 μl) of each of the three samples was harvested, using sterile syringes and re-sealing the empty wells, at the respective timepoint (2 time points, including exponential (EX) and stationary (ST)).

#### Measured growth rate estimation

All measured growth rates were estimated using R (v.4.1.3) with multiple packages for data wrangling (i.e., openxlsx, tidyr, reshape2, comprehenr) and statistical modeling (i.e., stats, GenSA, robustbase). Overall, the bacterial growth rates were estimated by the log ratio of the change in OD over time dlog_2_(OD)/dt.

The first batch of EX samples in *E. coli* (23MS06, 23MS10, 23MS13) was assessed by manual OD measurements in 30 min intervals (min. 7 time points until harvesting for fast-growing conditions). To reduce impacts of background OD and condition-dependent differences in lag phase length, a simulated annealing model was used to globally optimize the exponential fit:

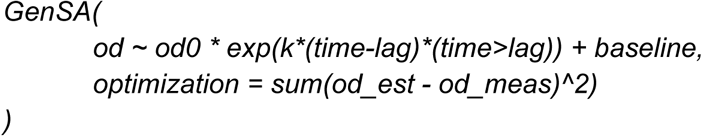

, where GenSA is the generalized simulated annealing function, od is the OD at the time point of harvesting, od0 is the starting OD at time point zero and k is the inferred exponential growth rate. Further, time is the amount of time until harvesting, lag is the amount of time before growth starts (during lag phase), the baseline is a constant added as a background OD (i.e., the mean OD during the first three time points across replicates). The function is optimized to minimize the sum of squared errors between estimated and measured OD.

All other experimental batches in *E. coli* (22MS23) and *B. theta*/*A. rectalis* (23MS10) used automated plate reader measurements in 10 min intervals across growth phases. Due to shorter intervals and more consistent measurements, the growth rates were estimated by a linear fit to log-scaled OD values including the time point of sample collection and six time points prior to collection thus based on a time interval of 60 mins:

lm( log(od) ∼ time, data=subset(time>=lag.sa) )

, where lm is the linear model function, log(od) are log-scaled OD values, time is the amount of time until harvesting, lag.sa is the lag phase duration extracted from the simulated annealing model and the data subset used for model fitting contained all time points after lag phase has ended thus when growth starts.

### DNA/RNA extraction from cultures

For all *E. coli* samples, the culture pellets were subjected to enzymatic lysis in 50 μl 1x Tris-EDTA containing 8 mg/ml Lysozyme for 10 min at room temperature (RT) and vortexed for 10 seconds every 2 minutes. Subsequently, 600 μl RLT buffer (Allprep DNA/RNA Mini kit, Qiagen) and one tube of 100 micron Zirconium beads (OPS Diagnostics LLC) were added to the cells. The cells were mechanically lysed 2 times at 30 Hz for 5 min (Retsch MM400 Mixer Mill, Fisher Scientific) with an incubation step at RT for 5 min in between. After the custom lysis protocol, DNA/RNA was extracted following the standard protocol of the Allprep DNA/RNA Mini extraction kit (Qiagen) with an on-column DNase digest (RNase-free DNase set, Qiagen) to prevent DNA contamination in the extracted RNA.

For all *B. theta* and *A. rectalis* culture pellets, DNA/RNA was extracted following the standard protocol of the Allprep Power Fecal Pro DNA/RNA extraction kit (Qiagen) without on-column DNase digest. The following volumes were adapted to maximize the yield due to the small harvested culture volume (250 μl): After adding CD2 buffer, 650 μl supernatant was transferred to a new tube and 650 μl of CD3 buffer was added. The DNA column was loaded two times with 650 μl and and to each flow-through, 325 μl of 96-100% ethanol was added. The RNA column was loaded three times with 650 μl.

After all DNA/RNA extractions, 10 μl 3M sodium acetate, 275 μl ice-cold 100% ethanol and 1 μl glycogen (5 mg/ml) were added to 100 μl DNA/RNA eluate for overnight ethanol precipitation at -20 °C. Ethanol precipitation consisted of initial centrifugation (30 min, 13000 rpm, 4°C) to create a pellet, supernatant removal, two wash steps of the pellet with 500 μl ice-cold 75% ethanol (by repeated centrifugation (10 min, 13000 rpm, 4°C) and supernatant removal) and redissolving in respective amounts of RNase-free water, yielding purified DNA (100 μl) and RNA

(50 μl). Quality controls for yield, purity and integrity were performed by Qubit (Qubit 3 Fluorometer, Invitrogen), Nanodrop (NanoDrop Eight, Labtech) and Fragment Analysis (Fragment Analyzer, Agilent) respectively. The DNA/RNA yields were maximized by harvesting the whole culture volume. The purity of DNA/RNA was targeted to be at a A260/280 ratio of 1.8 and 2.0 respectively and a A230/260 ratio of 2.0-2.2. The integrity was targeted to be at a DNA Integrity Number (DIN) >= 7.0 and RNA Integrity Number (RIN) >= 8.5 (max DIN/RIN = 10.0).

### DNA/RNA library preparation and sequencing

Depending on the obtained yield, 20-200 ng of purified DNA was sheared into 350 bp fragments (LE220-plus Focused-ultrasonicator, Covaris). To target this fragment size, the shearing time was optimized depending on the detected fragment size by Fragment Analyzer. Genomic DNA libraries with a target length of 500 bp were generated with the Ultra II DNA Library Prep Kit (NEBNext) from 10-500 ng sheared DNA (depending on experimental batch) and either using CleanNGS (LabGene) or Sera-Mag Select (Cytiva) beads with adjusted ratios.

We used 100-1000 ng of total RNA (depending on growth phases) to generate RNA libraries with an insert length of 190 bp and a target length of 450 bp with the Stranded Total RNA Prep Kit, Ligation with Ribo-Zero Plus (Illumina). Sera-Mag Select beads (Cytiva) were used for all cleanups except for the RNA cleanup before reverse transcription, in which RNAClean XP beads (Beckman Coulter) were used.

All multiplexed DNA/RNA libraries were evaluated in terms of yield and sizes by Fragment Analysis and pooled for sequencing. In case of observed primer dimers in the library pool, a repeated cleanup with Sera-Mag Select beads (0.7X) was performed. The final library pool was spiked with 3% PhiX to control for read diversity. The spiked library pool was sequenced on a NextSeq2000 platform (Illumina) at the Genome Engineering and Measurement Lab (GEML, https://geml.ethz.ch/) which is part of the Functional Genomics Center Zurich (FGCZ). A total of 400M-1.1B (P2/P3 kit) paired end (PE) reads (2x100/2x150, 200/300 Cycles) were sequenced to target 5M reads per sample.

### Primary processing of sequencing data

#### Quality control of raw genomic/transcriptomic reads

BBMap^70^ (v.38.71) was used for quality control of raw sequencing reads by removing adapters from the reads, removing reads mapping to quality control sequences (PhiX genome) and discarding low-quality reads (trimq = 14, maq = 20, maxns = 1, and minlength = 45). In addition, the T-overhangs in sequencing adapters used for rapid ligation, resulting in a “T” at the first position in all transcriptomic sequencing reads, were trimmed.

#### Mapping of read counts

The quality-controlled reads were aligned to each of the reference genomes of the three strains (<u>CP092639.1</u>, <u>CP092641.1</u>, <u>CP092643.1</u>) using BWA^71^ (v.0.7.17). The alignments were filtered using in-house software to be at least 45 bp in length, with an identity of ≥97% and a coverage

of ≥80% of the read sequence (i = 0.97, a = 45, c = 0.8) and converted into sorted bam files using SAMtools^72^ (v.1.17). The raw read counts per gene were obtained using featureCounts^73^ (v.2.0.3) with fractional counts for multi-mappers (-M --fraction) and counting read pairs (-p

--countReadPairs).

### Quality control of mapped transcriptomic reads per strain

To ensure sufficient quality of the sequencing data, we explored the sequencing depth and clustering between whole transcriptomes in each of the three EAM strains. After removing genes with low counts (averaged counts < 10 across samples), saturation curves were generated from subsampled read counts in the range [0.1,1] multiplied by the raw sequencing depth. While saturation of detected numbers of genes was observed in all samples for *E. coli* and *B. theta*, six *A. rectalis* samples (STAU23-2_23MS10_Ere_Glu_7-0_an_37_TP6_ST_1/2/3, STAU23-2_23MS10_Ere_Mal_7-0_an_37_TP6_ST_1/2/3) displayed a linear correlation of detected genes with subsampling with a maximum number of detected genes < 500 and were thus discarded (Supplementary Fig. 6; remaining = 34/40 samples).

### Transcriptomic signatures of bacteria growing in vitro

#### Differential expression across all genes

To assess growth phase-dependent global transcriptome shifts in each strain, we first filtered out genes with < 10 raw transcriptomic counts averaged across samples. Across all genes fulfilling this minimal expression requirement, we performed a differential expression analysis comparing EX to ST samples for each strain (*E. coli*: 87/30 EX/ST samples, *B. theta*: 36/34, *A. rectalis*: 18/9) using pyDESeq2^63^ (v.0.4.10). Genes were classified as differentially expressed according to adjusted p-value < 0.05, using Benjamini-Hochberg correction for multiple testing, and absolute log_2_-fold changes > 2. To evaluate the overrepresentation of functional groups in either up-/ or downregulated genes, we performed an enrichment analysis on COG categories assigned by eggNOG-mapper^74^ (v.2.1.12) in each strain, within either significantly upregulated (padj < 0.05; log_2_FC > 2) or downregulated (padj < 0.05; log_2_FC < -2) genes compared to all genes using Fisher’s exact test (with Benjamini Hochberg multiple testing correction). To test the enrichment of COG categories among the strongest differentially expressed genes, we generated a subset of the top 10% up/- or downregulated genes (according to log_2_FCs) and performed a Fisher’s exact test (with Benjamini Hochberg multiple testing correction) against all other differentially expressed genes.

#### Extraction of transcriptomic MG counts

With the aim to explore transcriptomic MG profiles independent of the availability of whole genomes, we extracted the raw transcriptomic counts of single-copy, phylogenetic marker genes. To this end, we ran GTDB-Tk^75^ (v.2.3.0) and in-house software on the three reference genomes to extract all marker gene sequences used within GTDB^45^ and proGenomes2^46^ respectively. Redundant sequences across the two databases were removed using VSEARCH (v.2.15.0) with exact match search (perc_id = 1.0), resulting in 129 unique marker genes

(Supplementary Table 3). 127 out of the 129 marker genes were identified in all three strains (*E. coli*, *B. theta* and *A. rectalis*) and thus used for downstream analyses. The raw counts of these 127 marker genes were extracted from the raw transcriptomic counts across all genes by creating a map between marker genes and genes by sequence alignment using VSEARCH (v.2.15.0, perc_id = 0.95).

#### Differential expression across MGs

To inspect differential expression in the 127 MGs, we subset our results from differential expression analysis across all genes to the MGs, adjusting the threshold for differential expression (adjusted p-value < 0.05; absolute log_2_-fold change > 1) to also consider less strong changes in expression levels. We determined the number of MGs with congruent differential expression across all strains, according to significant differential expression (adjusted p-values < 0.05 in all strains) and the same directionality (either log_2_FC > 1 or log_2_FC < -1 in all strains).

### Bacterial growth modeling in vitro

#### Transcriptomic growth phase classification

All transcriptomic-based growth modeling was carried out in Python (v.3.10.15) using the packages numpy (v.1.25.1) and pandas (v.2.0.3) for data wrangling, scikit-learn (v.1.3.0) for modeling and plotly (v.5.22.0) as well as seaborn (v.0.13.2) for visualization.

To develop a method purely dependent on raw transcriptomic counts of the 127 MGs, we used an internal approach for MG normalization by disregarding transcriptomic counts of all other genes. To obtain normalized MG counts, we performed gene length normalization by dividing raw read counts of each MG by its gene length (in kilobases), resulting in reads per kilobase (RPK). To normalize for sequencing depth within the MGs, we divided the RPK values by a scaling factor (i.e., the sum of RPK values across all MGs in each sample divided by one million), resulting in Transcripts Per Million MG counts (TPM_mg). Lastly, TPM_mg counts were log_2_-scaled after adding a pseudo-count of 0.5, resulting in transcriptomic MG profiles (log_2_(TPM_mg + 0.5)).

To explore transcriptomic signatures across growth phases, a Principal Component Analysis (PCA) with 20 PCs was performed on normalized transcriptomic counts of the 127 MGs in *E. coli* EX/TR/ST samples. Next, we trained a binary growth classifier (3 PCs, 9 kNNs) with 5x cross validation in *E. coli* EX/ST samples. For 5x cross validation, the samples were split into training/test data (0.8/0.2) ensuring replicates remain together and an even distribution of growth states across splits (sklearn.model_selection.StratifiedGroupKFold).

#### Application across species in vitro

We then applied the developed classifier to *E. coli* TR samples, *B. theta* and *A. rectalis* samples. The normalized transcriptomic MG counts were transformed to plot them on the first two PCs of a PCA on *E. coli* EX/ST samples. For inspection of misclassifications in *A. rectalis*, we assessed the hierarchical clustering (metric = euclidean, method = ward) of whole transcriptomes, according to log_2_(TPM) counts, of all *A. rectalis* EX/ST samples (Supplementary Fig. 3).

#### Robustness testing with missing marker genes

Since these single-copy, phylogenetic MGs are expected to be 77-95% universal according to GTDB^45^ (Release 220), and thus could be missing from some genomes, we tested the robustness of our *E. coli*-trained classifier in *B. theta* and *A. rectalis* by *in silico* pairwise removal of 2 out of the 127 marker genes from raw transcriptomic MG counts. We evaluated imputing missing marker genes with the mean of raw transcriptomic counts across the rest of the marker genes in each sample and found that they have negligible effects on precision and accuracy both in *B. theta* and *A. rectalis* (Supplementary Fig. 7).

#### Genomic growth rate prediction

To compare transcriptomic, marker gene-based growth prediction to existing methods, the genomic PTR-based tool CoPTR^29^ (v.1.0.0) was applied to all quality-controlled genomic reads from one end (_1.fq.gz) with default parameters. The required minimum sequencing depth to generate robust log_2_(PTR) estimates was by far exceeded with the targeted sequencing depth per strain (5M reads).

To evaluate potential biases in log_2_(PTR) estimates obtained for EX samples and identify sample-specific factors, such as cultivation conditions (i.e., temperature) or strains potentially driving these biases, a linear model (sklearn.linear_model.LinearRegression) was fit to all *E. coli* EX samples grown at 37 °C. The residuals to the linear model were calculated for different temperature ranges in *E. coli* and for different strains (*B. theta*, *A. rectalis*) at 37 °C.

### In vivo growth classification in EAM mice

#### Mouse experiments and cecum content samples

To investigate metagenomic/metatranscriptomic signatures of bacterial growth in a community setting *in vivo*, we leveraged cecum content samples from EAM (Easily Accessible Microbiota) mice, harboring the same three strains (*E. coli*, *B. theta* and *A. rectalis*) that were collected as a part of a different study^64^. All mouse experiments were from previously conducted experimental batches and were approved by the Swiss Kantonal authorities (License ZH120/19 and ZH016/21) according to the legal and ethical requirements. The mice were euthanized at different daytimes (2 pm, 5 pm, 9 pm) and wet cecum content was collected, immediately flash-frozen and stored at -80 °C. A detailed description of the EAM mice and mouse experiments can be found in the respective publication, however, the sample metadata are provided (Supplementary Table 4) and experimental processing is described below as the cecum samples were processed as a part of this study.

### Experimental sample processing

For all EAM mouse cecum content samples, DNA/RNA was extracted following the standard protocol of the Allprep Power Fecal Pro DNA/RNA extraction kit (Qiagen) without on-column DNase digest. 20-50 mg of flash-frozen cecum content was weighed into PowerBead Pro tubes with sterile spatulas before the lysis buffer was added. 5 μl RNase inhibitors (Superase-In RNase Inhibitor, Invitrogen) were added to the lysis buffer to prevent potential RNase activity. DNA/RNA purification by ethanol precipitation, quality control, library preparation and sequencing were conducted according to *in vitro* sample processing.

### Community composition by qPCR

To assess cell numbers per gram of dry weight of cecum content across daytimes, a subsample of approx. 20-50 mg cecum content was weighed to determine the wet weight, freeze-dried for 12 hours and weighed again to determine the dry weight. DNA/RNA was re-extracted from this subsample according to the description above. A qPCR was performed with 2 μl DNA template added to 6 μl of Ultra-Pure H_2_O, 10 μl qPCR Master Mix (FastStart Universal SYBR Green Master (Rox)) and 1 μl each of strain-specific forward/reverse primers (Supplementary Table 6) amounting to a total volume of 20 μl. In addition, DNA standards from known cell numbers per taxon were added to the qPCR plate to create standard curves and infer absolute cell numbers. The qPCR protocol included the following steps: hold stage with 50°C for 2 min and 95°C for 10 min, PCR stage with 40 cycles of 95°C for 15 sec and 60°C for 1 min and melt curve stage with 95°C for 15 sec, 60°C for 1 min and 95°C for 15 sec. Cell counts per μl of extracted DNA were quantified by flow cytometry prior to DNA extraction (*E. coli*: 2.9*10^7^, *B. theta*: 7.08*10^7^, *A. rectalis*: 3.54*10^7^ cells/μl of DNA). Based on the experimental qPCR protocol described in section 2.1.7 and using a dilution series of these DNA standards (undiluted, 1:10^1^, 1:10^2^, 1:10^3^, 1:10^4^, 1:10^5^, 1:10^6^) as templates, strain-specific standard curves were inferred. Absolute cell counts in the EAM cecum content samples were inferred from a linear fit between the quantified cell numbers of the standards and obtained qPCR cycle thresholds (Ct). Relative abundances were inferred from absolute cell counts to describe the community composition in the EAM mouse cecum across daytimes.

### Primary processing of sequencing data

Quality control of the metagenomic/metatranscriptomic sequencing data and the generation of normalized, transcriptomic marker gene counts was conducted according to *in vitro* data processing. For read mapping, the raw metatranscriptomic reads were mapped to a concatenated fasta file containing the three EAM bacterial genomes. Sample 2 (at 2 pm) was discarded due to low sample integrity (bloody cecum), large amounts of host contamination and distinctiveness from expected raw metagenomic/metatranscriptomic reads per strain.

### In vivo metatranscriptomic growth classification

The 127 MGs were extracted for all EAM strains as described in *in vitro* methods. Raw transcriptomic MG counts were internally normalized (described in *Methods – Transcriptomic growth phase classification*) and our *E. coli*-trained growth classification model was applied to the normalized MG counts. For further investigation, we performed hierarchical clustering (metric = euclidean, method = ward) of normalized transcriptomic counts across all genes (i.e., log_2_(TPM), rescaled to the range [0,1]).

### In vivo metatranscriptomic growth rate regression for E. coli

The large number of samples and wide range of measured growth rates across growth phases in *E. coli* were leveraged to go beyond growth phase classification and evaluate different machine learning-based regression models (from sklearn.linear_model and sklearn.ensemble: Ridge, Lasso, Elastic Net, Random Forest, Gradient Boosting, Ensemble) with 5x cross validation using all *E. coli* samples (EX/TR/ST). For 5x cross validation, the samples were split into training/test data (80/20%) ensuring replicates remain together and an even growth rate distribution across splits (sklearn.model_selection.StratifiedGroupKFold). Standard scaling was applied to normalized transcriptomic MG counts (sklearn.preprocessing.StandardScaler). The performance of the different models with selected parameter sets (Supplementary Table 5) across folds was evaluated by mean absolute error (MAE) and R-squared (sklearn.metrics.mean_absolute_error, .r2_score). The best-performing Ridge regression model was applied to predict growth rates of *E. coli* in EAM cecum samples *in vivo*.

### In vivo metagenomic growth rate regression

The genomic-based growth rate prediction tool CoPTR^29^ was applied to all quality-controlled genomic reads from one end (_1.fq.gz) to generate log_2_(PTR) estimates for each reference genome across all samples. The minimum sequencing depth required to generate robust log_2_(PTR) estimates^29^ was exceeded with the targeted sequencing depth per sample (60M reads), resulting in an approximate sequencing depth of 6M reads for the lowest abundant member *E. coli*, based on its expected relative abundance (10%) and similar genome sizes (*E. coli*: 4.6*10^6^ bp, *B. theta*: 6.3*10^6^ bp, *A. rectalis*: 3.4*10^6^ bp).

### In vivo growth classification in Oligo mice

The previously conducted study^43^ investigated effects of intravenous lipopolysaccharide (LPS) injection on systemic and intestinal inflammation in mice, perturbations of their gut microbiota and opportunities for pathogen blooms. The study applied intravenous LPS injection and PBS controls in 6 mice each and euthanized the mice 6h post injection to obtain cecum samples and generate metatranscriptomic data. Any detailed description of the methods from experimental design to generating raw metatranscriptomic counts is available in the publication.

### Marker gene extraction and imputation of missing marker gene counts

The 127 MGs were extracted for the three Oligo-MM^12^ bacterial genomes of interest (*Enterocloster clostridioformis* YL32: <u>CP015399.2</u>, *Blautia coccoides* YL58: <u>CP015405.2</u>, *Bacteroides caecimuris* I48: <u>CP015401.2</u>) as described in *in vitro* methods. Most of the MGs were identified in the three strains apart from one missing marker gene in *E. clostridioformis* YL32 (COG0100) and one missing marker gene in *B. caecimuris* I48 (TIGR00020). Raw

metatranscriptomic MG counts were thus extracted from raw metatranscriptomic counts across all genes and unidentified marker genes were imputed by mean metatranscriptomic counts across the remaining marker genes.

### In vivo metatranscriptomic growth classification

Internal normalization (described in *Methods - Transcriptomic growth phase classification*) was applied to obtain normalized transcriptomic MG counts (log_2_(TPM_mg + 0.5)). The *E. coli*-trained growth classifier was applied to the normalized transcriptomic counts to obtain growth classifications across treatments (LPS, PBS). Statistical significance between the treatment groups was assessed by running a Mann-Whitney U test without correction for multiple testing (as the number of tests was small).

## Contributions

M.L.S. and S.S. conceptualized and designed the study. M.L.S. conducted the *in vitro* cultivation experiments and processed all *in vitro* EAM strain samples and *in vivo* EAM cecum content samples for sequencing. G.G. and M.A. provided the *in vivo* EAM cecum samples across daytimes. M.L.S. conducted the formal analysis and investigation. M.L.S., A.S., M.D. and S.M.V. contributed to method development. M.L.S., H.J.R. and A.S. developed the software. M.L.S. visualized results and wrote the manuscript draft and A.S., S.S., M.D., S.M.V., W.D.H., M.A. and E.S. contributed to review and editing. A.S. and S.S. supervised the study. E.S., W.D.H. and S.S. contributed to funding acquisition.

## Acknowledgements

This work was funded by the NCCR Microbiomes, a National Centre of Competence in Research, provided by the Swiss National Science Foundation to S.S., E.S. and W.D.H. (51NF40_180575 and 51NF40_225148). S.S. and E.S. were supported by the Basel Research Centre for Child Health Multi-Investigator Project 2020 (BRCCH_MIP: Microbiota Engineering for Child Health). S.M.V. acknowledges funding from the Human Frontier Science Program (HFSP) through a postdoctoral fellowship [LT0050/2023-L]. The authors acknowledge the Genome Engineering and Measurement Lab (GEML, https://geml.ethz.ch/), which is part of the Functional Genomics Center Zurich (FGCZ), for providing walk-in access and services for short-read (meta)genomic and (meta)transcriptomic sequencing. The authors would like to acknowledge the support of the IT Service and HPC facilities of ETH Zürich. The authors thank Sanne Kroon for providing access to the metatranscriptomic data from Oligo-MM^12^ cecum content samples, Guillem Salazar for methodological advice at early stages of the project and Alessio Milanese for providing access to a code basis for marker gene extraction.

## Supplementary Figures

**Supplementary Figure 1.**
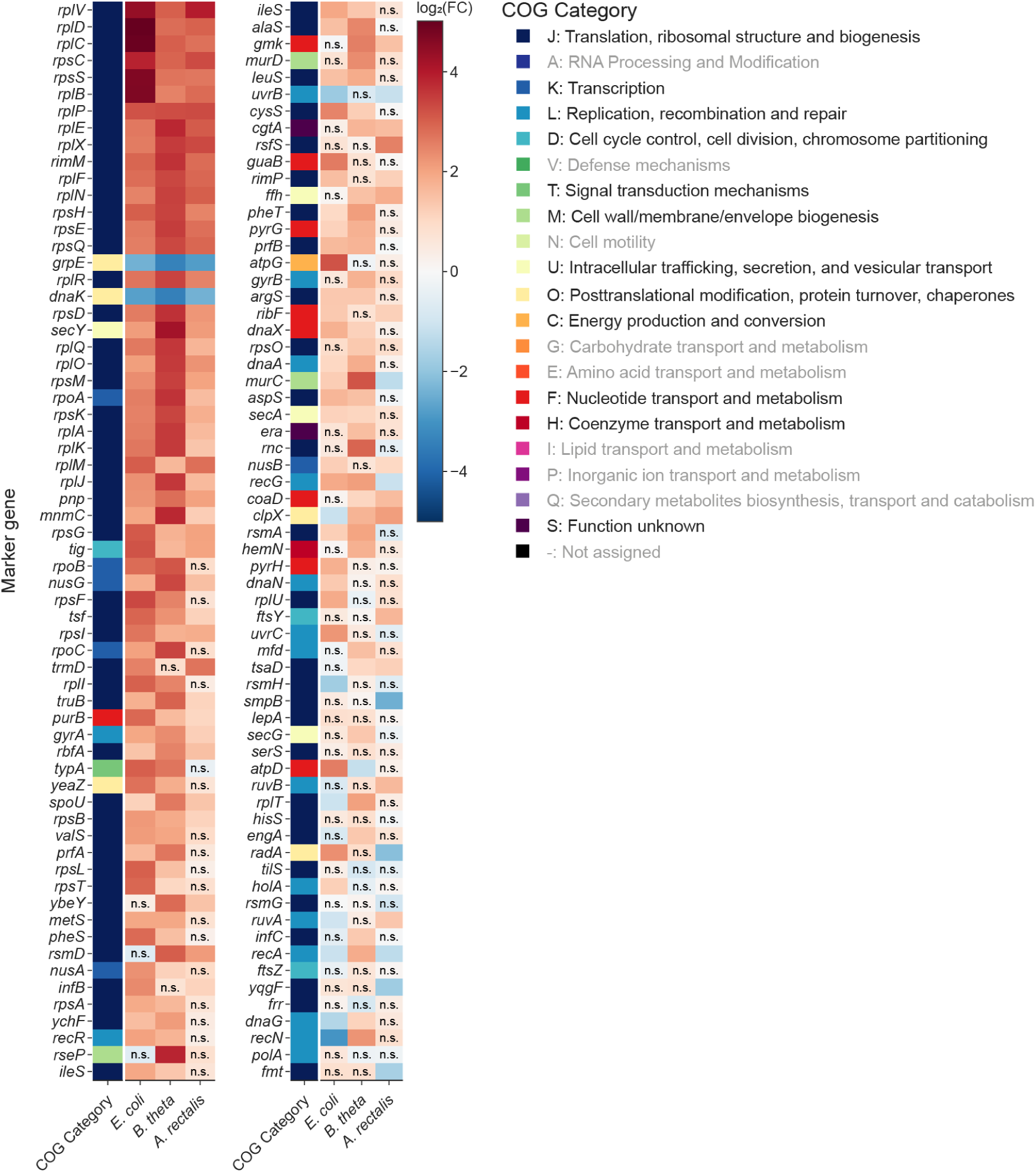
L**o**g2 **fold changes (log_2_FC) of all 127 MGs based on differential expression in each of the three strains.** The MGs were labelled according to assigned COG categories, compared across strains and sorted by mean log_2_FC across strains. Significant differential expression was evaluated for each MG in each strain (padj < 0.05; absolute log_2_FC > 1), resulting in non-significant (n.s.) labels.

**Supplementary Figure 2.**
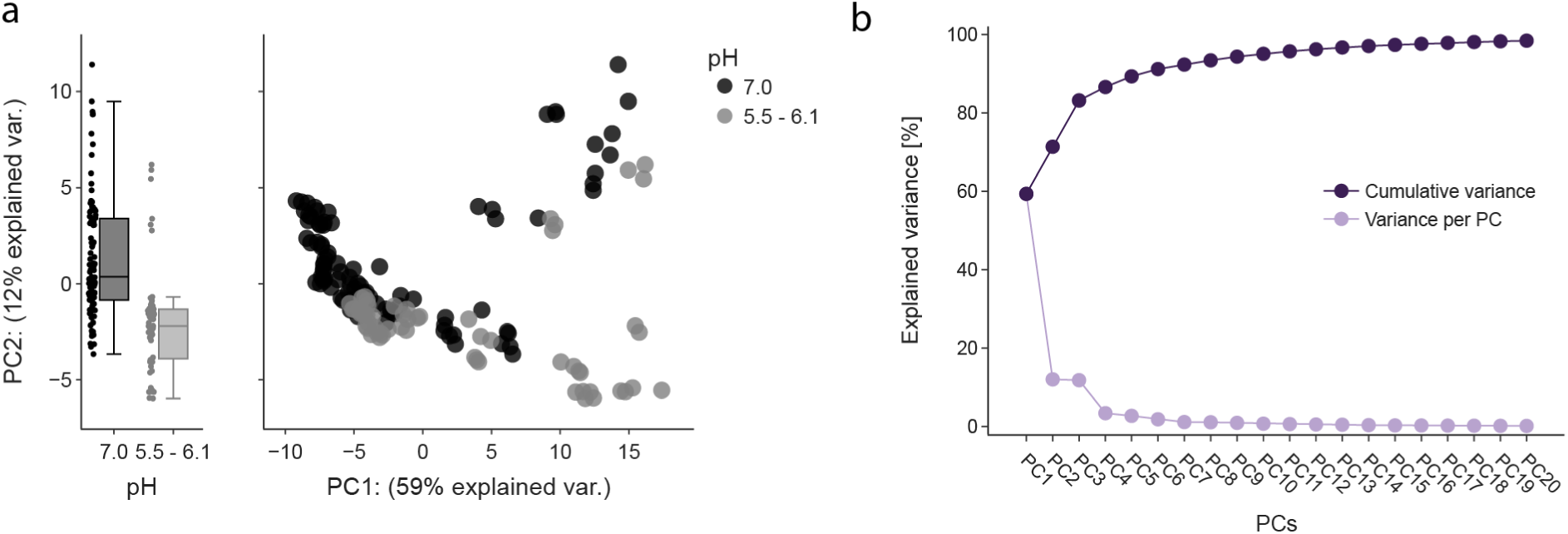
V**a**riation **in other PCs beyond PC1. a,** A PCA was performed on all E. coli samples (EX/TR/ST samples shown in Fig. 2b) and yielded a partial separation by pH along PC2, particularly in ST phase samples (right side on PC1). **b,** The explained variance for 1-20 PCs (light purple: per PC, dark purple: cumulative) was assessed from a PCA on all *E. coli* samples (EX, TR, ST).

**Supplementary Figure 3.**
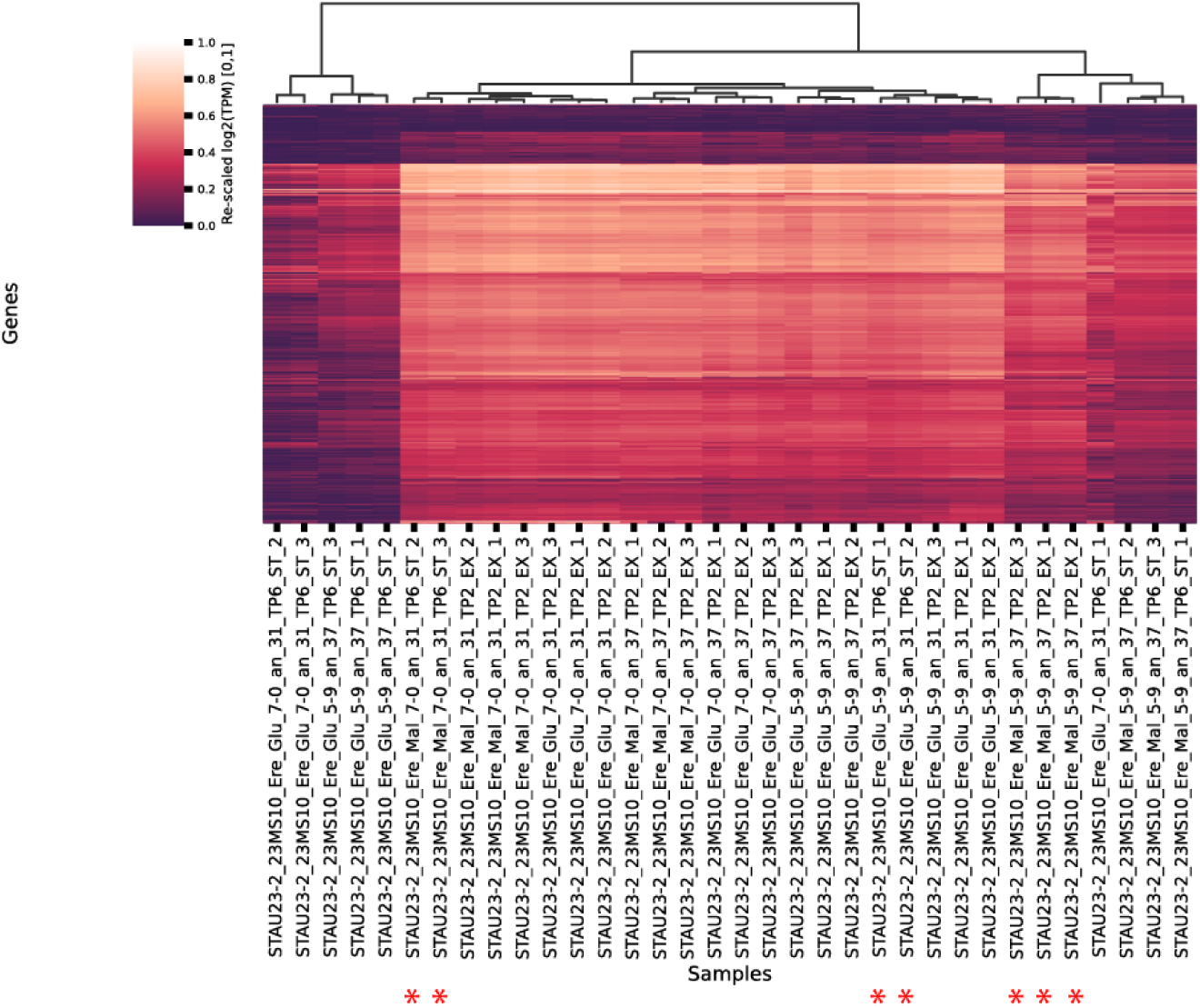
W**h**ole **transcriptome clustering of *A. rectalis* samples.** Seven *A. rectalis* samples contradicted their experimentally assigned growth phases based on hierarchical clustering of the whole transcriptomes (i.e., log(TPM) across all genes, metric = euclidean, method = ward). The seven samples with inconclusive growth phase assignment are highlighted with red stars.

**Supplementary Figure 4.**
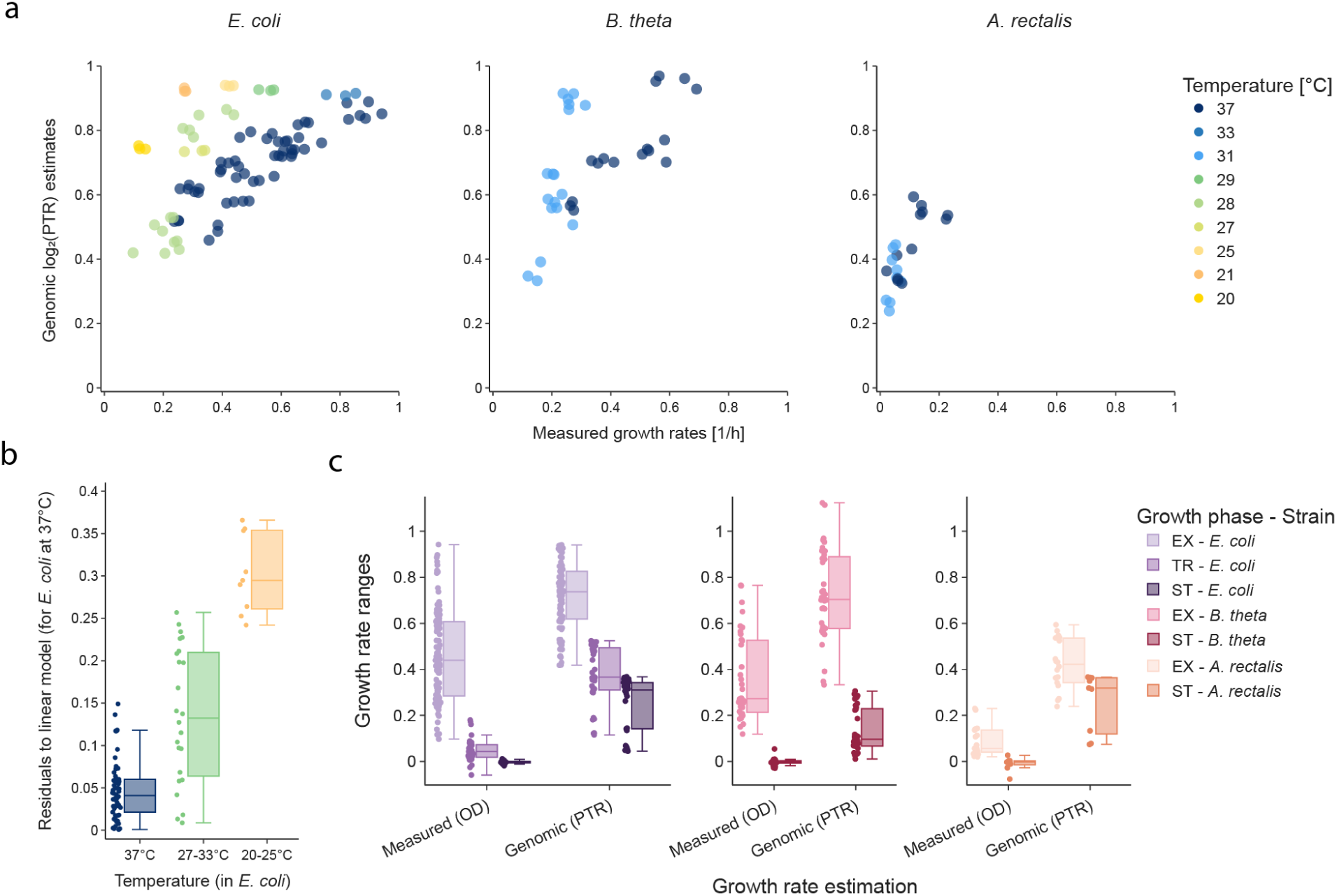
C**o**mparison **to existing methods for bacterial growth rate prediction using genomic data. a**, Correlation of measured growth rates (OD-based) versus genomic log_2_(PTR) estimates in exponential (EX) samples across the strains *E. coli* (left), *B. theta* (middle) and *A. rectalis* (right). **b**, Residuals to a linear model fitted to all exponential (EX) *E. coli* samples at 37 °C cultivation temperature across three temperature bins 37 ° (blue), 27-33 °C (green) and 20-25 °C (yellow). **c,** Ranges of measured growth rates (OD-based) compared to predicted genomic growth rate estimates (PTR-based) across growth phases in exponential/transition/stationary (EX/TR/ST) samples in *E. coli* (purple), *B. theta* (red) and *A. rectalis* (orange).

**Supplementary Figure 5.**
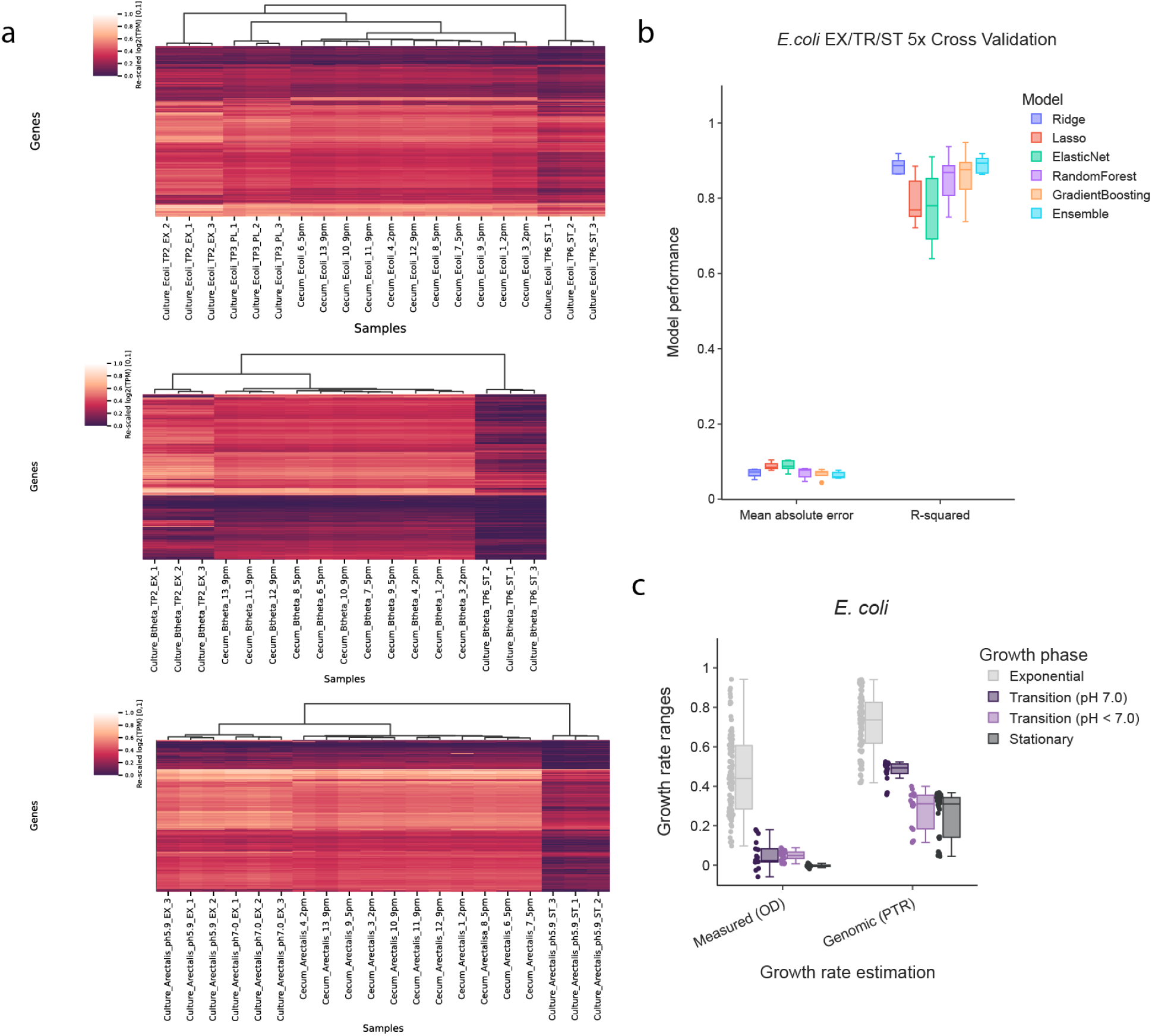
W**h**ole **transcriptome analyses supporting growth classification across growth phases. a,** Whole transcriptomes (i.e., log_2_(TPM) scaled to the range of [0,1]) of *in vivo* samples were clustered together with a subset of *in vitro* reference samples across growth phases in each strain (*E. coli*: 22MS23-Glucose-pH7-37°C EX/PL/ST samples; with PL samples being the first TR sample during a plateau of net zero growth reached right after exiting EX phase, *B. theta*: 23MS10-Glucose-pH7.0-37°C EX/ST samples, *A. rectalis*: 23MS10-Glucose-pH7/5.9-37°C EX/ST samples). These samples were chosen according to their expected highest similarity to *in vivo* conditions in the EAM mouse cecum. **b**, We evaluated the model performance of different transcriptomic, marker gene-based regression models for growth rate prediction with 5x cross validation in *E. coli* by Mean Absolute Error (MAE) and R-Squared. **c**, We compared *E. coli in vivo* log_2_(PTR) estimates from EAM cecum content to *in vitro* estimates at a comparable pH (7.0) from *E. coli* TR samples.

**Supplementary Figure 6.**
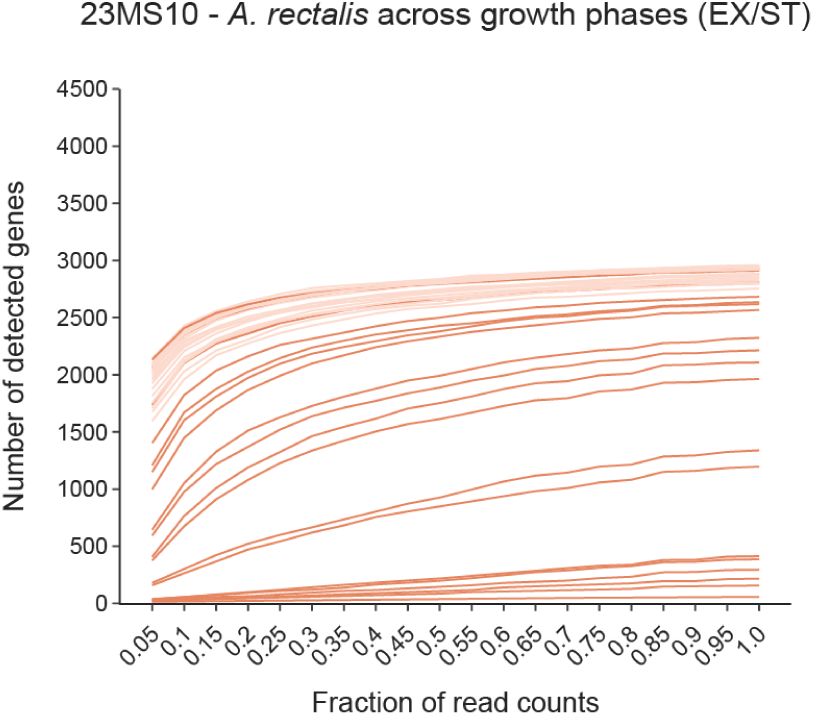
D**i**scarded ***A. rectalis* samples due to insufficient sample and/or sequencing data quality. a,** Six *A. rectalis* samples displayed a linearly increasing correlation between the number of detected genes across subsampled sequencing depth in the range of [0.1,1.0] of the original sequencing depth, with a maximum number of detected genes < 500.

**Supplementary Figure 7.**
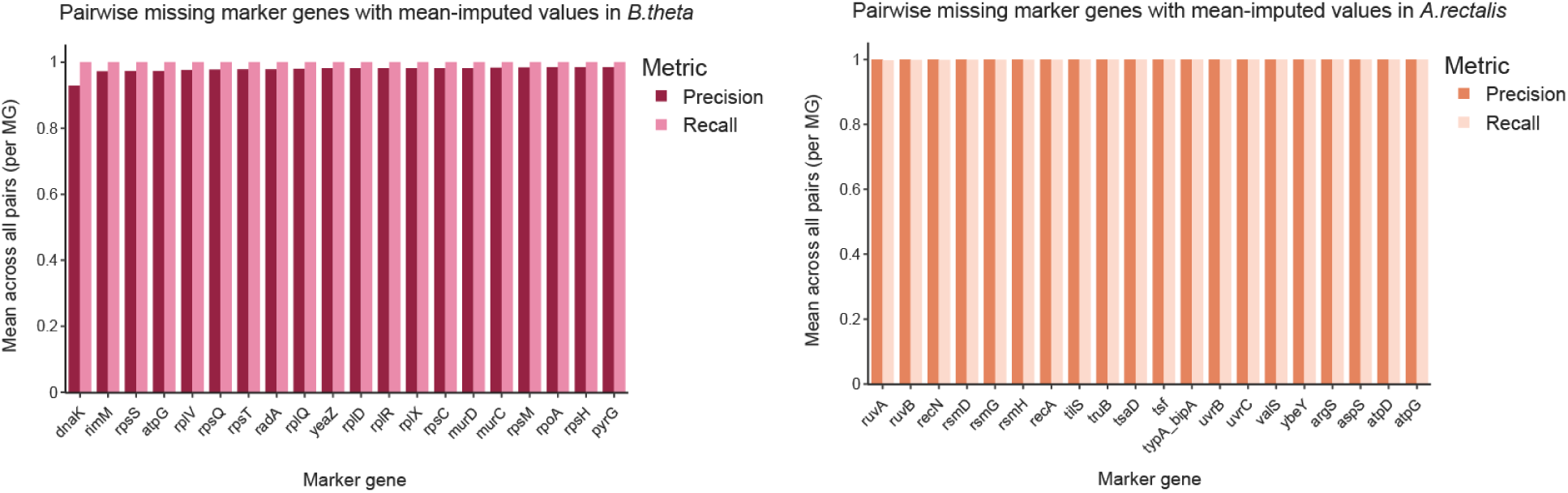
E**f**fects **of pairwise removal and imputation of missing marker genes on classifier performance *in vitro*.** Pairs of MGs were removed from the *B. theta*/*A. rectalis* raw transcriptomic MG counts and imputed by the mean raw transcriptomic counts across all remaining MGs in each sample. After imputation of missing marker genes, log_2_(TPM_mg + 0.5) normalization was performed and the performance (precision/recall) of the *E. coli*-trained classifier was tested in predicting growth states in *B. theta*/*A. rectalis*. The MGs were sorted in descending order according to the mean effect on performance across one MG paired with all other MGs for removal and imputation.

## Supplementary Tables

**Supplementary Table 1.** Cultivation conditions across bacterial strains and experimental batches. Abbreviations: EX = Exponential, PL = Plateau, IH = Inhibition, SG = Slow growth, ST = Stationary.

| Strain | Batch | Base medium | Carbon source | pH | Temperature | Timepoints |
| --- | --- | --- | --- | --- | --- | --- |
| *E. coli* | 21MS06 | M9 + Wolin's TE | Arabinose, Fructose, Gluconate, Glucose, Maltose, Succinate | 7.0 | 37 | EX |
|  |  |  | Ribose | 7.0 | 37, 27, 20 | EX |
|  | 21MS10 | M9 + Wolin's TE | Glucose | 7.0, 6.1, 5.9, 5.7, 5.5 | 37 | EX |
|  | 21MS13 | M9 + Wolin's TE | Glucose | 7.0 | 37, 33, 29, 25, 21 | EX |
|  | 22MS23 | M9 + Wolin's TE | Glucose | 7.0 | 37 | EX1, EX2, PL, IH, SG, ST1, ST2 |
|  |  |  |  | 5.5 | 37 | EX, PL, ST |
|  |  |  |  | 7.0 | 28 | EX, PL, ST |
|  |  |  |  | 5.5 | 28 | EX, PL, ST |
|  |  |  | Maltose | 7.0 | 37 | EX, PL, ST |
|  |  |  |  | 5.5 | 37 | EX, PL, ST |
|  |  |  |  | 7.0 | 28 | EX, PL, ST |
|  |  |  |  | 5.5 | 28 | EX1, EX2, PL, SG1, SG2, ST1, ST2 |
| *B. theta* | 23MS10 | Adapted TYG | Glucose | 7.0, 5.9 | 37 | EX, ST |
|  |  |  |  |  | 31 | EX, ST |
|  |  |  | Maltose | 7.0, 5.9 | 37 | EX, ST |
|  |  |  |  |  | 31 | EX, ST |
|  |  |  | Ribose | 7.0, 5.9 | 37 | EX, ST |
|  |  |  |  |  | 31 | EX, ST |
| *A. rectalis* | 23MS10 | Adapted TYG | Glucose | 7.0, 5.9 | 37 | EX, ST |
|  |  |  |  |  | 31 | EX, ST |
|  |  |  | Maltose | 7.0, 5.9 | 37 | EX, ST |
|  |  |  |  | 7.0 | 31 | EX, ST |

**Supplementary Table 2.** COG Enrichment in either all or top 10% up-/ or downregulated genes in the three EAM strains. The enrichment values are shown as n_enr/n_all (padj), with n_enr = number of genes in the enrichment group, n_all = number of genes in background used for comparison, and padj = adjusted p-value after multiple testing correction. COG enrichment analysis was first conducted for testing enrichment in either up/downregulated genes against all genes as a background (column 3 and 4). Secondly, COG enrichment analysis was performed on top 10% up/downregulated genes in comparison to all differentially expressed (DE) genes (column 5 and 6).

| COG Category | Strain | Enriched in Up vs All | Enriched in Down vs All | Enriched in Top10% Up vs DE | Enriched in Top 10% Down vs DE |
| --- | --- | --- | --- | --- | --- |
| J | *E. coli* | 70/240 (8.14e-16) | 8/240 (1.00e+00) | 16/78 (2.39e-02) | 2/78 (1.00e+00) |
| A | *E. coli* | 0/3 (1.00e+00) | 1/3 (9.62e-01) | 0/1 (1.00e+00) | 0/1 (1.00e+00) |
| K | *E. coli* | 15/345 (1.00e+00) | 79/345 (2.90e-01) | 2/94 (1.00e+00) | 10/94 (1.00e+00) |
| L | *E. coli* | 5/196 (1.00e+00) | 22/196 (1.00e+00) | 1/27 (1.00e+00) | 0/27 (1.00e+00) |
| D | *E. coli* | 3/49 (1.00e+00) | 4/49 (1.00e+00) | 2/7 (5.42e-01) | 1/7 (1.00e+00) |
| V | *E. coli* | 1/36 (1.00e+00) | 6/36 (1.00e+00) | 0/7 (1.00e+00) | 1/7 (1.00e+00) |
| T | *E. coli* | 11/115 (1.00e+00) | 30/115 (2.90e-01) | 3/41 (1.00e+00) | 6/41 (1.00e+00) |
| M | *E. coli* | 20/263 (1.00e+00) | 40/263 (1.00e+00) | 3/60 (1.00e+00) | 1/60 (1.00e+00) |
| N | *E. coli* | 21/107 (1.46e-02) | 22/107 (9.62e-01) | 10/43 (4.01e-02) | 4/43 (1.00e+00) |
| W | *E. coli* | 0/1 (1.00e+00) | 0/1 (1.00e+00) | 0/0 (1.00e+00) | 0/0 (1.00e+00) |
| U | *E. coli* | 9/163 (1.00e+00) | 39/163 (3.59e-01) | 2/48 (1.00e+00) | 6/48 (1.00e+00) |
| O | *E. coli* | 11/122 (1.00e+00) | 18/122 (1.00e+00) | 4/29 (7.93e-01) | 5/29 (1.00e+00) |
| C | *E. coli* | 37/285 (3.03e-01) | 57/285 (9.62e-01) | 7/94 (1.00e+00) | 10/94 (1.00e+00) |
| G | *E. coli* | 13/253 (1.00e+00) | 70/253 (9.92e-03) | 5/83 (1.00e+00) | 3/83 (1.00e+00) |
| E | *E. coli* | 65/293 (6.02e-09) | 45/293 (1.00e+00) | 22/110 (1.37e-02) | 8/110 (1.00e+00) |
| F | *E. coli* | 53/234 (9.02e-08) | 26/234 (1.00e+00) | 16/79 (2.39e-02) | 6/79 (1.00e+00) |
| H | *E. coli* | 21/134 (1.47e-01) | 11/134 (1.00e+00) | 5/32 (6.48e-01) | 1/32 (1.00e+00) |
| I | *E. coli* | 4/105 (1.00e+00) | 25/105 (5.23e-01) | 1/29 (1.00e+00) | 10/29 (5.66e-03) |
| P | *E. coli* | 42/335 (3.25e-01) | 62/335 (1.00e+00) | 16/104 (1.96e-01) | 10/104 (1.00e+00) |
| Q | *E. coli* | 7/62 (1.00e+00) | 13/62 (9.62e-01) | 3/20 (7.93e-01) | 2/20 (1.00e+00) |
| S | *E. coli* | 22/773 (1.00e+00) | 224/773 (4.65e-11) | 7/246 (1.00e+00) | 37/246 (3.29e-02) |
| - | *E. coli* | 4/77 (1.00e+00) | 21/77 (2.90e-01) | 0/25 (1.00e+00) | 2/25 (1.00e+00) |
| J | *B. theta* | 64/167 (5.56e-10) | 6/167 (1.00e+00) | 15/70 (5.38e-02) | 1/70 (1.00e+00) |
| A | *B. theta* | 0/1 (1.00e+00) | 0/1 (1.00e+00) | 0/0 (1.00e+00) | 0/0 (1.00e+00) |
| K | *B. theta* | 20/241 (1.00e+00) | 28/241 (1.00e+00) | 4/48 (1.00e+00) | 7/48 (1.00e+00) |
| L | *B. theta* | 29/231 (1.00e+00) | 20/231 (1.00e+00) | 2/49 (1.00e+00) | 2/49 (1.00e+00) |
| D | *B. theta* | 10/38 (3.16e-01) | 1/38 (1.00e+00) | 2/11 (1.00e+00) | 0/11 (1.00e+00) |
| V | *B. theta* | 18/83 (4.54e-01) | 9/83 (1.00e+00) | 2/27 (1.00e+00) | 2/27 (1.00e+00) |
| T | *B. theta* | 15/180 (1.00e+00) | 37/180 (2.59e-02) | 0/52 (1.00e+00) | 10/52 (2.99e-01) |
| M | *B. theta* | 84/332 (7.05e-04) | 35/332 (1.00e+00) | 17/119 (5.17e-01) | 12/119 (1.00e+00) |
| N | *B. theta* | 2/21 (1.00e+00) | 1/21 (1.00e+00) | 0/3 (1.00e+00) | 0/3 (1.00e+00) |
| W | *B. theta* |  |  |  |  |
| U | *B. theta* | 11/73 (1.00e+00) | 8/73 (1.00e+00) | 2/19 (1.00e+00) | 2/19 (1.00e+00) |
| O | *B. theta* | 12/88 (1.00e+00) | 20/88 (4.35e-02) | 4/32 (1.00e+00) | 2/32 (1.00e+00) |
| C | *B. theta* | 53/192 (1.13e-03) | 22/192 (1.00e+00) | 12/75 (5.17e-01) | 4/75 (1.00e+00) |
| G | *B. theta* | 52/364 (1.00e+00) | 42/364 (1.00e+00) | 12/94 (1.00e+00) | 10/94 (1.00e+00) |
| E | *B. theta* | 41/196 (3.16e-01) | 22/196 (1.00e+00) | 8/63 (1.00e+00) | 6/63 (1.00e+00) |
| F | *B. theta* | 24/107 (3.16e-01) | 12/107 (1.00e+00) | 4/36 (1.00e+00) | 2/36 (1.00e+00) |
| H | *B. theta* | 31/172 (9.86e-01) | 17/172 (1.00e+00) | 2/48 (1.00e+00) | 6/48 (1.00e+00) |
| I | *B. theta* | 17/67 (3.05e-01) | 7/67 (1.00e+00) | 3/24 (1.00e+00) | 4/24 (1.00e+00) |
| P | *B. theta* | 34/297 (1.00e+00) | 57/297 (2.30e-02) | 6/91 (1.00e+00) | 11/91 (1.00e+00) |
| Q | *B. theta* | 2/26 (1.00e+00) | 2/26 (1.00e+00) | 0/4 (1.00e+00) | 0/4 (1.00e+00) |
| S | *B. theta* | 143/899 (1.00e+00) | 133/899 (1.71e-01) | 20/276 (1.00e+00) | 27/276 (1.00e+00) |
| - | *B. theta* | 22/193 (1.00e+00) | 37/193 (4.35e-02) | 5/59 (1.00e+00) | 12/59 (2.27e-01) |
| J | *A. rectalis* | 30/172 (1.37e-09) | 3/172 (1.00e+00) | 17/33 (3.43e-09) | 0/33 (1.00e+00) |
| A | *A. rectalis* | 7/245 (1.00e+00) | 46/245 (2.32e-03) |  |  |
| K | *A. rectalis* | 6/198 (1.00e+00) | 28/198 (5.09e-01) | 2/53 (1.00e+00) | 2/53 (1.00e+00) |
| L | *A. rectalis* | 2/57 (1.00e+00) | 6/57 (1.00e+00) | 1/34 (1.00e+00) | 3/34 (1.00e+00) |
| D | *A. rectalis* | 2/93 (1.00e+00) | 7/93 (1.00e+00) | 0/8 (1.00e+00) | 4/8 (9.42e-02) |
| V | *A. rectalis* | 7/172 (1.00e+00) | 12/172 (1.00e+00) | 2/9 (1.00e+00) | 0/9 (1.00e+00) |
| T | *A. rectalis* | 10/145 (4.79e-01) | 9/145 (1.00e+00) | 3/19 (1.00e+00) | 1/19 (1.00e+00) |
| M | *A. rectalis* | 3/81 (1.00e+00) | 8/81 (1.00e+00) | 1/19 (1.00e+00) | 0/19 (1.00e+00) |
| N | *A. rectalis* | 0/1 (1.00e+00) | 0/1 (1.00e+00) | 1/11 (1.00e+00) | 0/11 (1.00e+00) |
| W | *A. rectalis* |  |  |  |  |
| U | *A. rectalis* | 5/55 (4.79e-01) | 6/55 (1.00e+00) | 1/11 (1.00e+00) | 2/11 (1.00e+00) |
| O | *A. rectalis* | 4/77 (1.00e+00) | 19/77 (4.63e-03) | 0/23 (1.00e+00) | 6/23 (1.34e-01) |
| C | *A. rectalis* | 8/114 (4.83e-01) | 14/114 (1.00e+00) | 3/22 (1.00e+00) | 1/22 (1.00e+00) |
| G | *A. rectalis* | 3/167 (1.00e+00) | 18/167 (1.00e+00) | 0/21 (1.00e+00) | 1/21 (1.00e+00) |
| E | *A. rectalis* | 2/138 (1.00e+00) | 5/138 (1.00e+00) | 1/7 (1.00e+00) | 1/7 (1.00e+00) |
| F | *A. rectalis* | 2/79 (1.00e+00) | 3/79 (1.00e+00) | 0/5 (1.00e+00) | 0/5 (1.00e+00) |
| H | *A. rectalis* | 3/114 (1.00e+00) | 7/114 (1.00e+00) | 1/10 (1.00e+00) | 1/10 (1.00e+00) |
| I | *A. rectalis* | 7/58 (1.79e-01) | 1/58 (1.00e+00) | 2/8 (1.00e+00) | 0/8 (1.00e+00) |
| P | *A. rectalis* | 8/98 (4.79e-01) | 16/98 (4.29e-01) | 3/24 (1.00e+00) | 1/24 (1.00e+00) |
| Q | *A. rectalis* | 2/14 (4.79e-01) | 0/14 (1.00e+00) | 1/2 (1.00e+00) | 0/2 (1.00e+00) |
| S | *A. rectalis* | 15/535 (1.00e+00) | 62/535 (1.00e+00) | 5/77 (1.00e+00) | 11/77 (6.31e-01) |
| - | *A. rectalis* | 6/211 (1.00e+00) | 50/211 (1.83e-06) | 1/56 (1.00e+00) | 11/56 (1.34e-01) |

**Supplementary Table 3.**
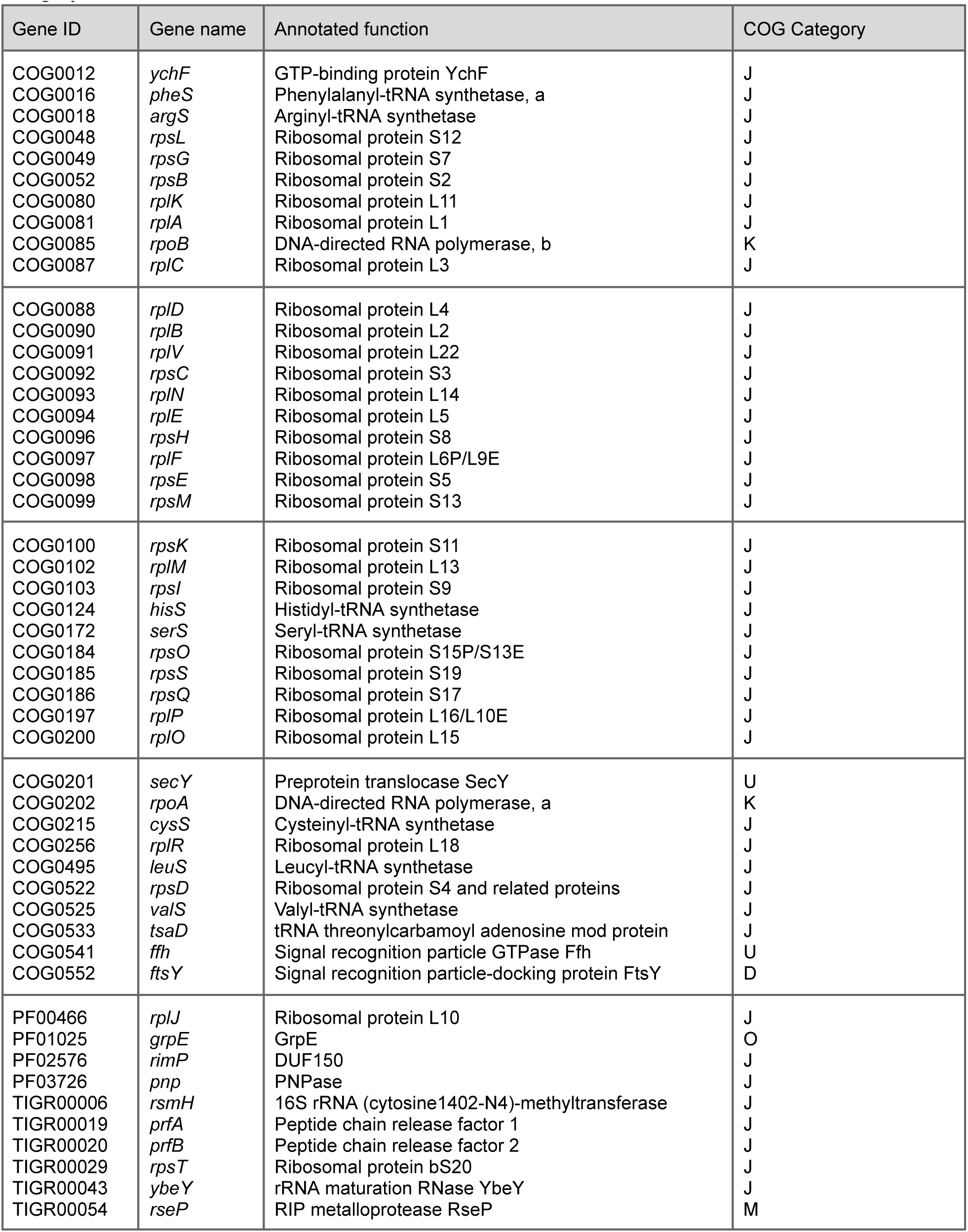
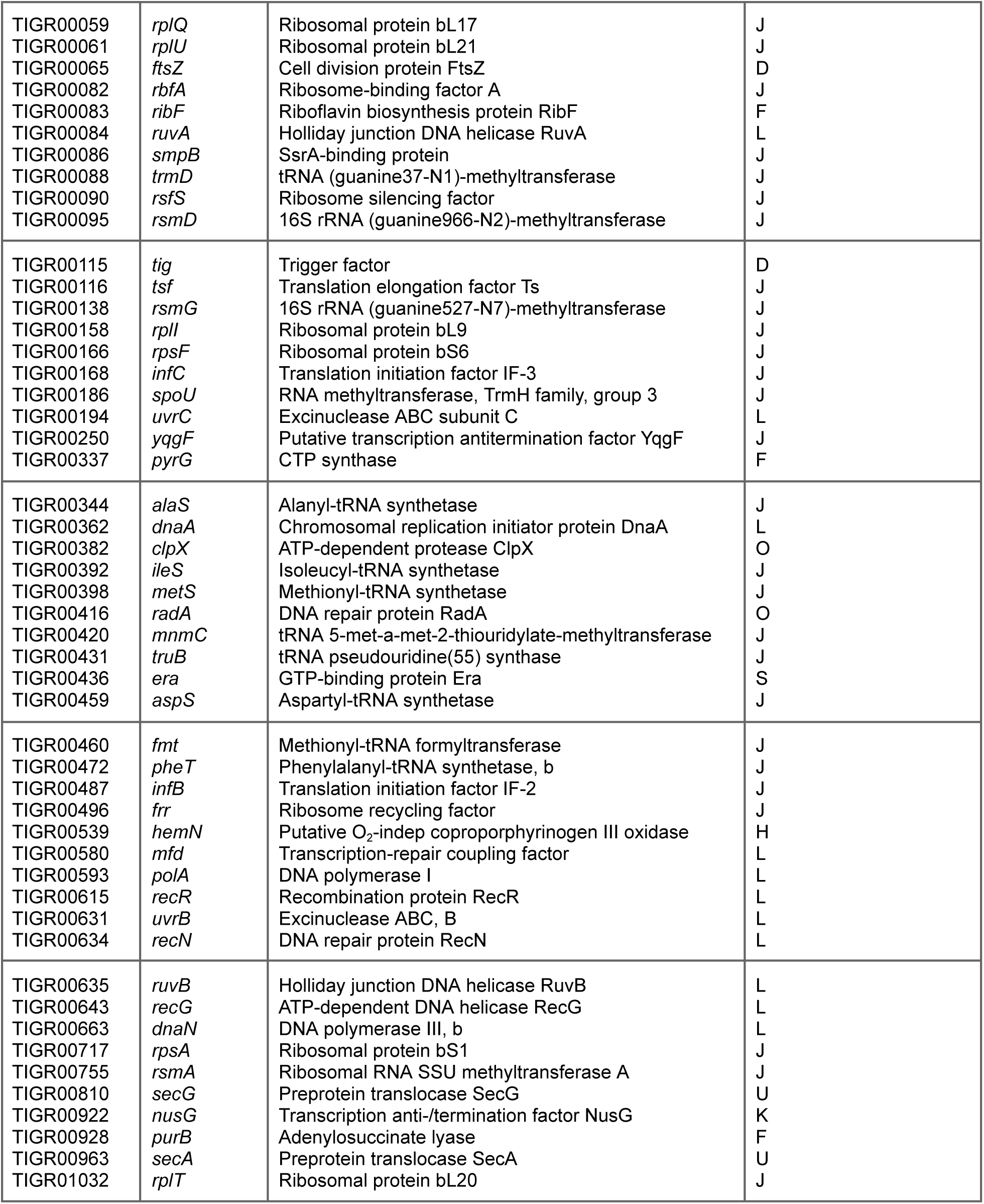
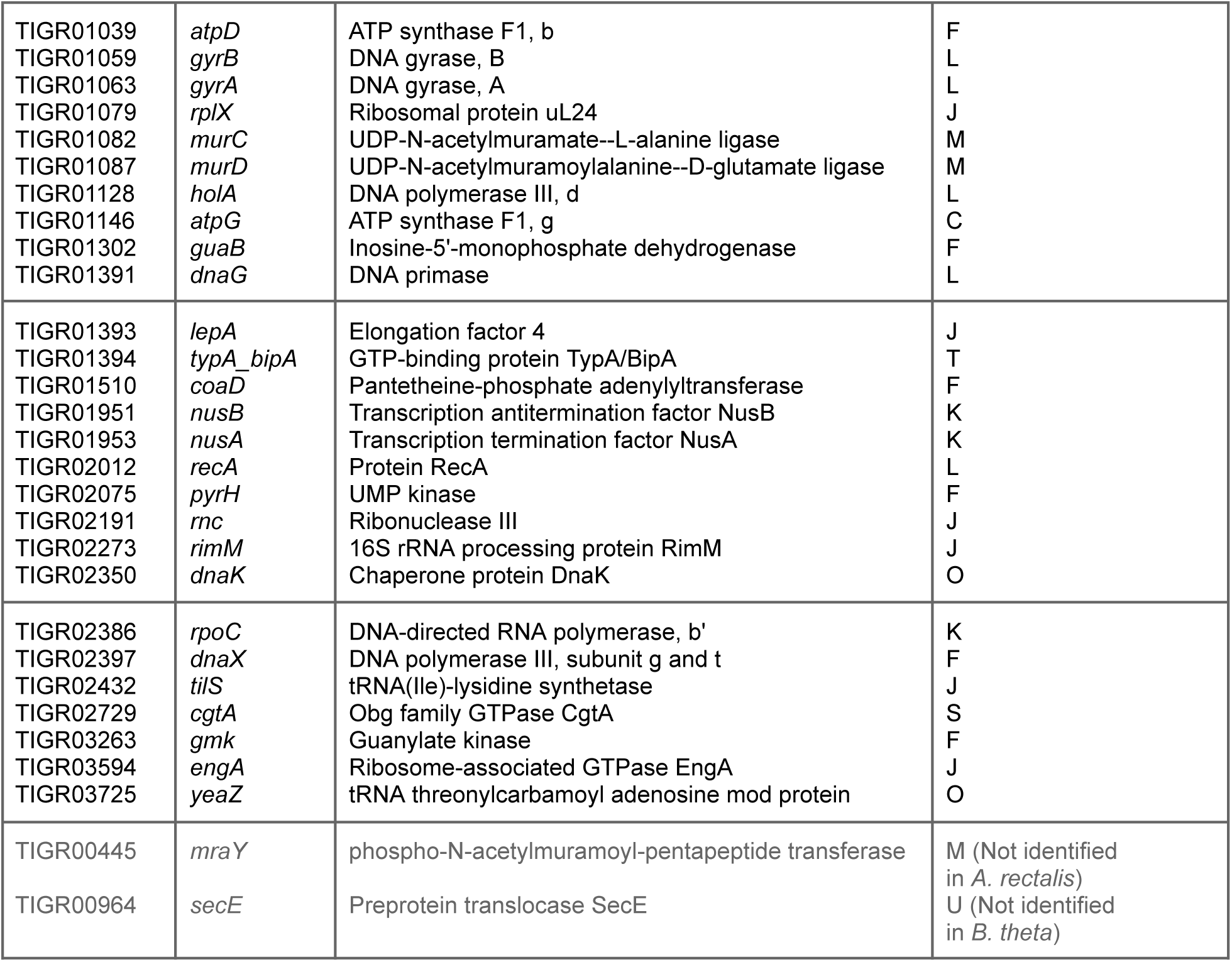
Overview of 129 marker genes, sorted alphabetically and displayed in chunks of size 10. For each marker gene, gene ID, gene name, annotated function and assigned COG category are listed.

**Supplementary Table 4.**
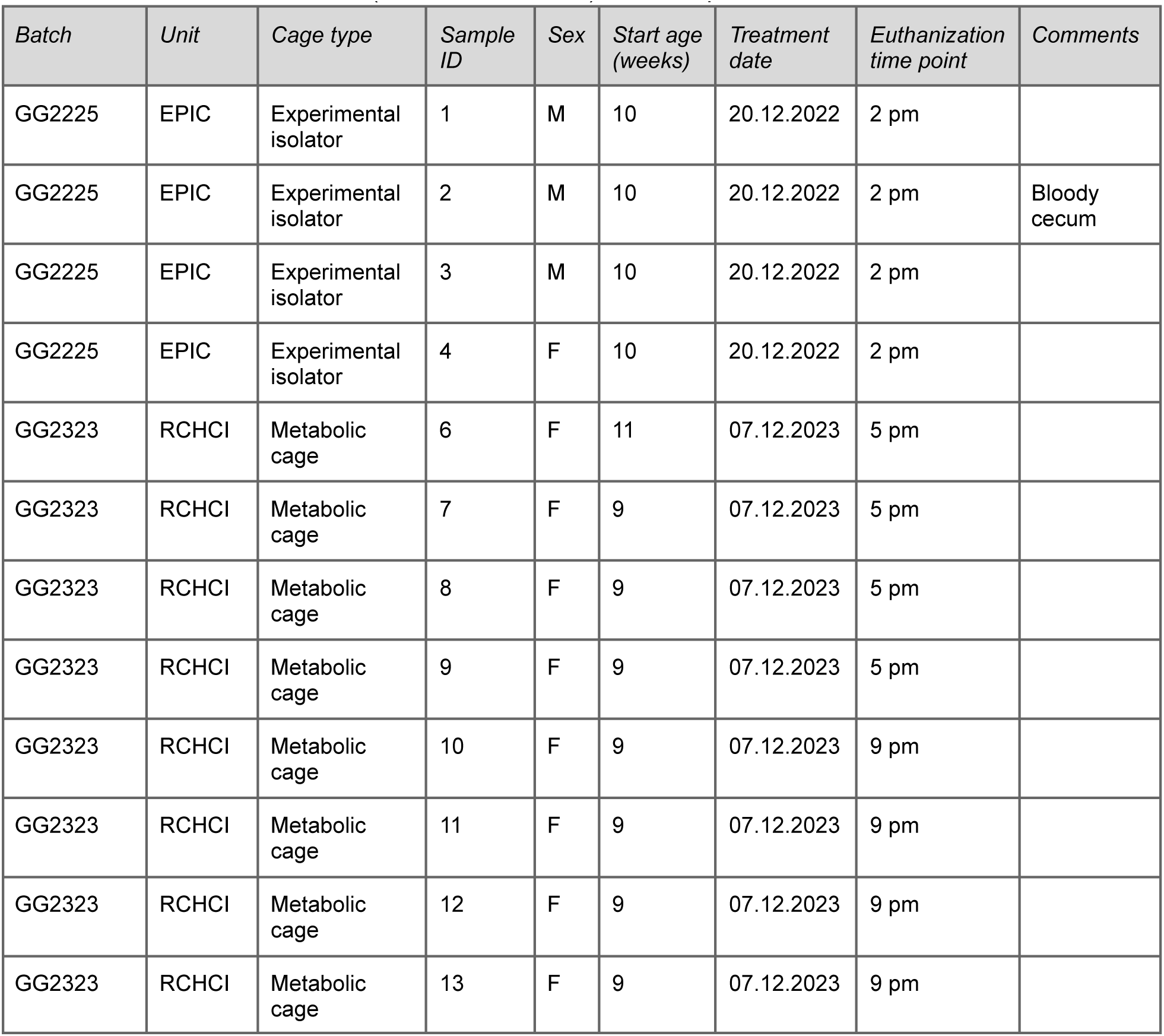
Metadata of the cecum content samples from EAM mice. The mice came from the same mouse colony, were all bred in the EPIC facility (ETH Phenomics Center) and transferred to another unit in EPIC or RCHCl (Rodent Center HCI) for the experiment.

| *Batch* | *Unit* | *Cage type* | *Sample ID* | *Sex* | *Start age (weeks)* | *Treatment date* | *Euthanization time point* | *Comments* |
| --- | --- | --- | --- | --- | --- | --- | --- | --- |
| GG2225 | EPIC | Experimental isolator | 1 | M | 10 | 20.12.2022 | 2 pm |  |
| GG2225 | EPIC | Experimental isolator | 2 | M | 10 | 20.12.2022 | 2 pm | Bloody cecum |
| GG2225 | EPIC | Experimental isolator | 3 | M | 10 | 20.12.2022 | 2 pm |  |
| GG2225 | EPIC | Experimental isolator | 4 | F | 10 | 20.12.2022 | 2 pm |  |
| GG2323 | RCHCI | Metabolic cage | 6 | F | 11 | 07.12.2023 | 5 pm |  |
| GG2323 | RCHCI | Metabolic cage | 7 | F | 9 | 07.12.2023 | 5 pm |  |
| GG2323 | RCHCI | Metabolic cage | 8 | F | 9 | 07.12.2023 | 5 pm |  |
| GG2323 | RCHCI | Metabolic cage | 9 | F | 9 | 07.12.2023 | 5 pm |  |
| GG2323 | RCHCI | Metabolic cage | 10 | F | 9 | 07.12.2023 | 9 pm |  |
| GG2323 | RCHCI | Metabolic cage | 11 | F | 9 | 07.12.2023 | 9 pm |  |
| GG2323 | RCHCI | Metabolic cage | 12 | F | 9 | 07.12.2023 | 9 pm |  |
| GG2323 | RCHCI | Metabolic cage | 13 | F | 9 | 07.12.2023 | 9 pm |  |

**Supplementary Table 5.** Machine learning-based regression models with tested parameter sets. Multiple regressors were evaluated with 5x Cross Validation in *E. coli* EX/TR/ST samples according to Mean Absolute Error (MAE) and R-Squared.

| *Model name* | *Final parameter choices* |
| --- | --- |
| Ridge | alpha = 10, fit_intercept = True |
| Lasso | alpha = 0.001, fit_intercept = True, max_iter = 3000, selection = "random" |
| Elastic Net | alpha = 0.001, l1_ratio = 0.5, max_iter = 10000 |
| Random Forest | max_features = 0.2, min_samples_split = 2, min_samples_leaf = 1, n_estimators = 100 |
| Gradient Boosting | max_features = 0.2, min_samples_split = 2, min_samples_leaf = 1, max_depth = 10, learning_rate = 0.1, subsample = 0.9 |
| Ensemble | VotingRegressor(estimators = [Ridge, Lasso, ElasticNet, RandomForest, GradientBoosting]) |

**Supplementary Table 6.**
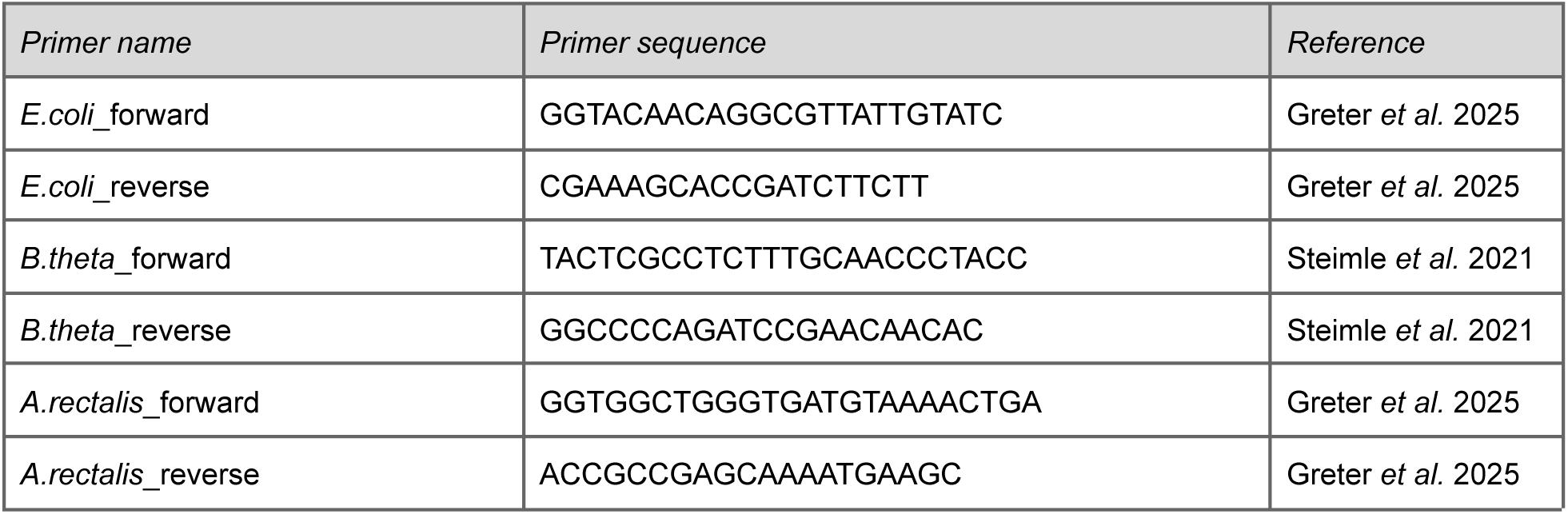
Strain-specific qPCR Primers for each of the three EAM strains.

